# RtmR is a membrane-embedded RRM-family RNA-binding protein

**DOI:** 10.1101/2025.07.10.664275

**Authors:** Jacob A. Vander Griend, Denise A. Ludvik, Hunter C. Nottage, Andrew Mehle, Mark J. Mandel

## Abstract

The animal symbiont *Vibrio fischeri* has served as a model organism for molecular processes underlying bacterial group behaviors, including quorum sensing and biofilm development. Here, using a genetic approach to identify negative regulators of biofilm formation in *V. fischeri*, we identified a membrane-bound RNA-binding protein, RtmR (VF_2432), that acts as an inhibitor of the symbiosis polysaccharide (SYP) biofilm. Membrane localization of the protein seems to be required for protein stability or expression, as truncation of the transmembrane helices led to an inability to detect the protein. The conserved RNP1 and RNP2 motifs in RtmR’s cytoplasmic RNA recognition motif (RRM) domain are required for function, and we demonstrate binding to RNA substrates. Identification of RtmR RNA ligands was conducted with a iiCLIP-seq approach that revealed a large interactome. One transcript identified was that of the biofilm regulatory histidine kinase RscS. We found that RtmR biofilm inhibition depends on RscS activity and that RtmR negatively regulates levels of RscS. Overall, this work characterizes a novel type of bacterial RNA-binding protein.

**IMPORTANCE:** Bacterial RNA-binding proteins (RBPs) perform key functions to regulate stress responses and development. Bacterial RBPs including the RNA chaperones Hfq and ProQ, the global regulator CsrA, and the cold shock proteins (Csps) have been extensively studied, although additional classes of RBPs have been predicted by bioinformatic methods including those carrying an RRM domain. This work expands on recent studies of RRM domain proteins in bacteria to characterize a membrane-bound RRM protein that regulates bacterial biofilm development. Given our rapidly-expanding knowledge regarding the role for RNA-binding proteins in bacterial molecular biology, this work contributes a new class of membrane-bound regulators with homologs in human pathogens and marine symbionts.

## INTRODUCTION

Biofilms, multicellular aggregates encased in secreted exopolysaccharides, are likely to be the predominant state for bacterial life (1). For those bacteria that colonize niches on or within eukaryotic hosts, biofilm formation provides benefits ranging from evasion of antimicrobials and host defenses to ensuring association with specific host tissues (2, 3). The symbiosis between the marine bioluminescent bacterium *Vibrio fischeri* and its animal host the Hawaiian bobtail squid (*Euprymna scolopes*) provides an opportunity to study the regulatory networks that control biofilm formation during natural colonization of an animal host. The biofilm formed from the symbiosis polysaccharide (SYP) is an essential component of the early stage of host colonization. *V. fischeri* strains that do not have the capability to produce SYP upon arrival in the host do not complete the colonization process, and regulation of SYP is a critical feature determining host colonization specificity (4–9). *V. fischeri* is also amenable to multiple genetic and molecular techniques that enable identification of regulators of symbiotic colonization (10–14).

In the early stages of squid colonization *V. fischeri* cells are captured from the surrounding seawater in a mucosal layer on the exterior of the light organ, a specialized symbiotic organ where the *V. fischeri* population is sequestered (15). *V. fischeri* cells entrained in this mucosal layer form biofilm-like aggregates as an essential first step in the colonization process, with aggregate cohesion dependent on the production of SYP (5, 6). SYP production is tightly regulated by *V. fischeri*, with a phosphorelay controlling the transcription of the 18-gene *syp* locus (16, 17). In strain ES114 the hybrid sensor kinase RscS initiates the SYP phosphorelay through phosphorylating another hybrid sensor kinase, SypF, which in turn phosphorylates response regulator SypG to activate *syp* expression (16, 18, 19). In opposition to RscS’s role as an activator of *syp* expression, BinK likely acts at the level of SypF to inhibit the SYP phosphorelay when environmental conditions are improper for SYP production (20, 21). Analysis of the Syp regulatory pathway has been made, in large part, through the use of genetic models carrying overexpression or deletion alleles of pathway members. One such model, *rscS\**, enables *in vitro* biofilm formation at the normal growth temperature of *V. fischeri* of 25 °C through the chromosomal overexpression of *rscS* from its native locus. A key biofilm phenotype of this model is the formation of highly wrinkled colony biofilms, which are dependent on an intact *syp* locus. Following earlier reports in the literature, we determined that growth of *rscS\** at the elevated growth temperature of 28 °C failed to yield the biofilm up phenotype observed in colony biofilms at the permissive temperature of 25 °C. In screening for explanatory factors for this phenotype we identified RtmR (VF_2432), a previously uncharacterized inner membrane-embedded RNA-binding protein, as an inhibitor of the symbiotic biofilm.

## RESULTS

### A genetic screen identifies RtmR (VF_2432) as a biofilm inhibitor

In culture, expression of the *syp* exopolysaccharide locus is barely detectable in *V. fischeri* squid symbiotic strain ES114 (5). Therefore, we used overexpression of *rscS* (i.e., *rscS\**), an established model of *in vitro* biofilm formation that stimulates *syp* locus expression through the same pathway used in host colonization (6, 22). This model produces wrinkled colony biofilms at 25 °C and colonies that are largely smooth at 28 °C (20). We generated a transposon insertion library in the *rscS\** background and screened the library to identify mutants with increased colony wrinkling at 28 °C. One mutant that enabled biofilm formation at 28 °C was a transposon insertion in *binK*, which encodes a known biofilm inhibitor (**Fig. 1A**) (20, 21). We additionally identified multiple biofilm-up mutants with unique insertions in and upstream of *VF_2432,* a gene with no previous association with SYP regulation, and that encodes an RNA recognition motif (RRM) domain protein. Below we characterize VF_2432 as an <u>R</u>RM-trans<u>m</u>embrane regulator and refer to it hereafter as RtmR.

**Figure 1.**
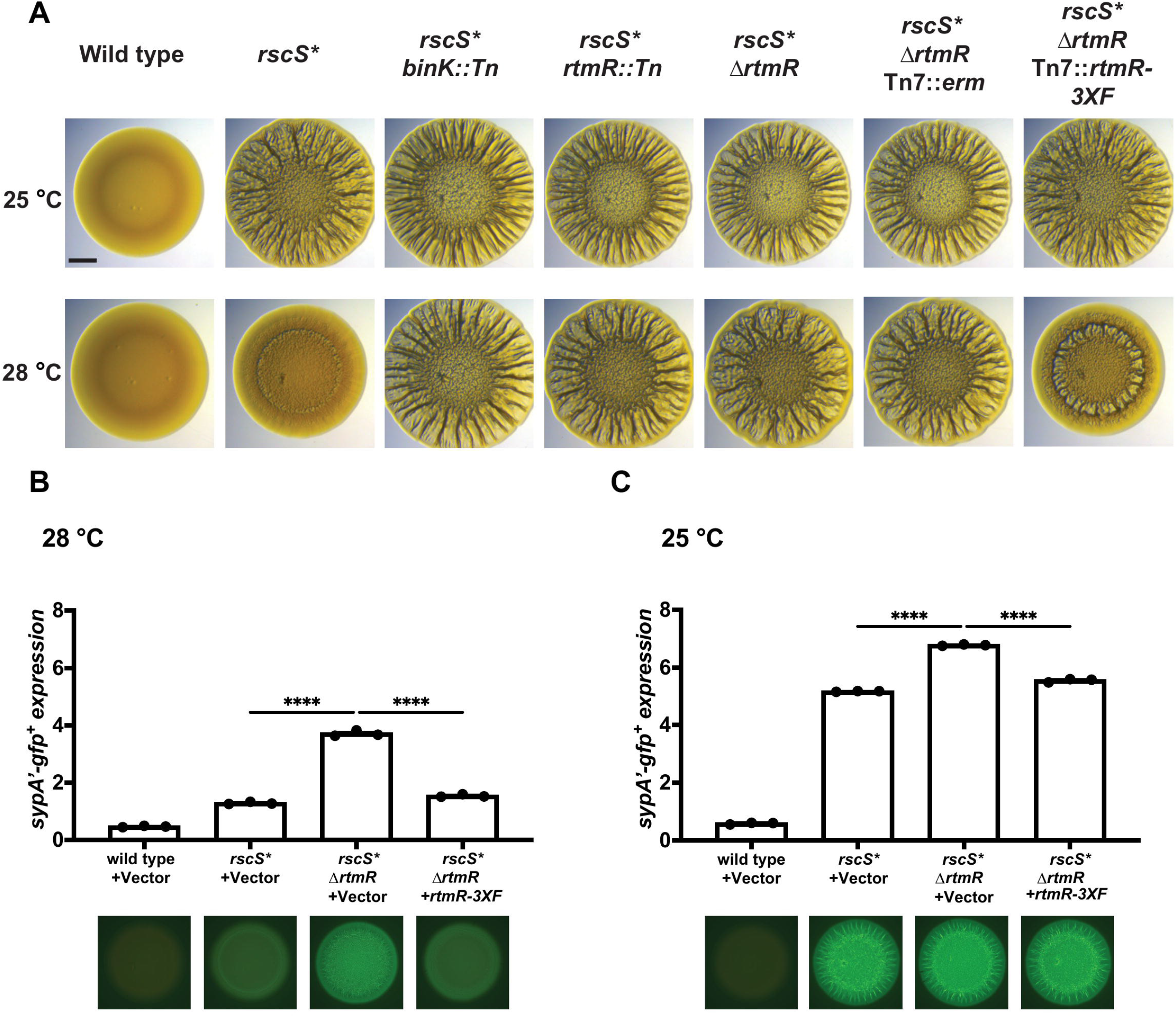
A genetic screen for biofilm inhibitors identifies RNA binding protein RtmR. A, Cultures of *V. fischeri* ES114 and indicated mutants were spotted on LBS agar and allowed to grow at 25 °C or 28 °C. Under these conditions, 25 °C is a biofilm permissive condition for the rscS* model and 28 °C is a biofilm restrictive condition. The *binK* and *rtmR* mutants that displayed wrinkled colony morphology are shown. At 48 h images of each strain were acquired using a Leica M60 microscope. All complementations were inserted at the neutral chromosomal *att*Tn*7* site. Scale bar is 2 mm. B, The indicated strains carrying a plasmid-based *sypA’-gfp*^+^ reporter were spotted on nutrient agar and allowed to grow at 28 °C for 24 h. Reporter activity was detected in each colony biofilm using a Zeiss Axio.Zoom macroscope, with *sypA’-gfp*^+^ expression normalized to a reporter backbone constitutive mCherry control. The + symbol indicates complementation at the *att*Tn*7* site with a vector control or *rtmR* allele. A total of n = 3 biological replicates were conducted, reported here as the average of two technical replicates. An ordinary one-way ANOVA was used to assess statistical significance of differences in *sypA* expression between the strains using Prism 10, with Šídák’s multiple comparison test. ****, P ≤ 0.0001. C, Analysis as in B but at 25 °C.

To validate that the gene product of *rtmR* was contributing to reduced biofilm formation at 28 °C, we generated an in-frame deletion of *rtmR* in the *rscS\** genetic background and examined the resulting strain’s capacity for biofilm formation. The deletion construct removed residues 2-119, 74% of the predicted 160-amino acid protein, so as to not impact the partially overlapping *murI* open reading frame. While *rscS\** and *rscS\** Δ*rtmR* both developed biofilms at 25 °C, the *rscS\** Δ*rtmR* strain recapitulated the transposon mutant phenotype of increased biofilm formation at 28 °C compared to *rscS\** alone. Complementation with a wild-type allele of *rtmR* reduced the colony biofilm, demonstrating that the phenotype was due to RtmR (**Fig. 1A**). In the *rscS\** strain, wrinkled colony formation is dependent on SYP production, so we hypothesized that RtmR may regulate *syp* expression (23). To test the hypothesis that RtmR was regulating *syp*, we used a plasmid-based transcriptional reporter of *sypA* (20, 24). At 28 °C, expression of *sypA* was higher in the *rscS\** Δ*rtmR* strain than in the *rscS\** or complemented strains (**Fig. 1B**). While this phenotype was most evident at 28 °C, we were also able to identify a milder increase in *sypA* expression at 25 °C as well (**Fig. 1C**), suggesting that RtmR is not a high-temperature specific regulator. Indeed, expression of RtmR appeared to be independent of growth temperature, as indicated by immunoblotting 3XFLAG tagged-RtmR from cultures grown at 25 °C and 28 °C (**Fig. S1**). Together, our results identify RtmR as an inhibitor of *V. fischeri* biofilm formation and *syp* transcription.

### RtmR activity is distinguishable from that of BinK in the *rscS\** biofilm model

Our initial data for RtmR paralleled that for the known biofilm inhibitor BinK. Both factors downregulate wrinkled colony formation at 28 °C, and both reduce *syp* locus transcription at 25 °C and 28 °C (20). To ask whether the two proteins act in the same pathway (e.g., whether RtmR regulation of *syp* was dependent on BinK), we measured *sypA* expression in deletion mutants of either *rtmR* or *binK,* and a double deletion of both inhibitors, with all assays in the *rscS\** parent (**Fig. 2**). At 28 °C, both single mutants robustly stimulated *syp* transcription compared to the parent strain, with Δ*binK* exhibiting a stronger phenotype than Δ*rtmR*. However, *sypA* expression in the double mutant was higher than either regulator mutant alone, suggesting that RtmR and BinK biofilm inhibition do not require the other’s function.

**Figure 2.**
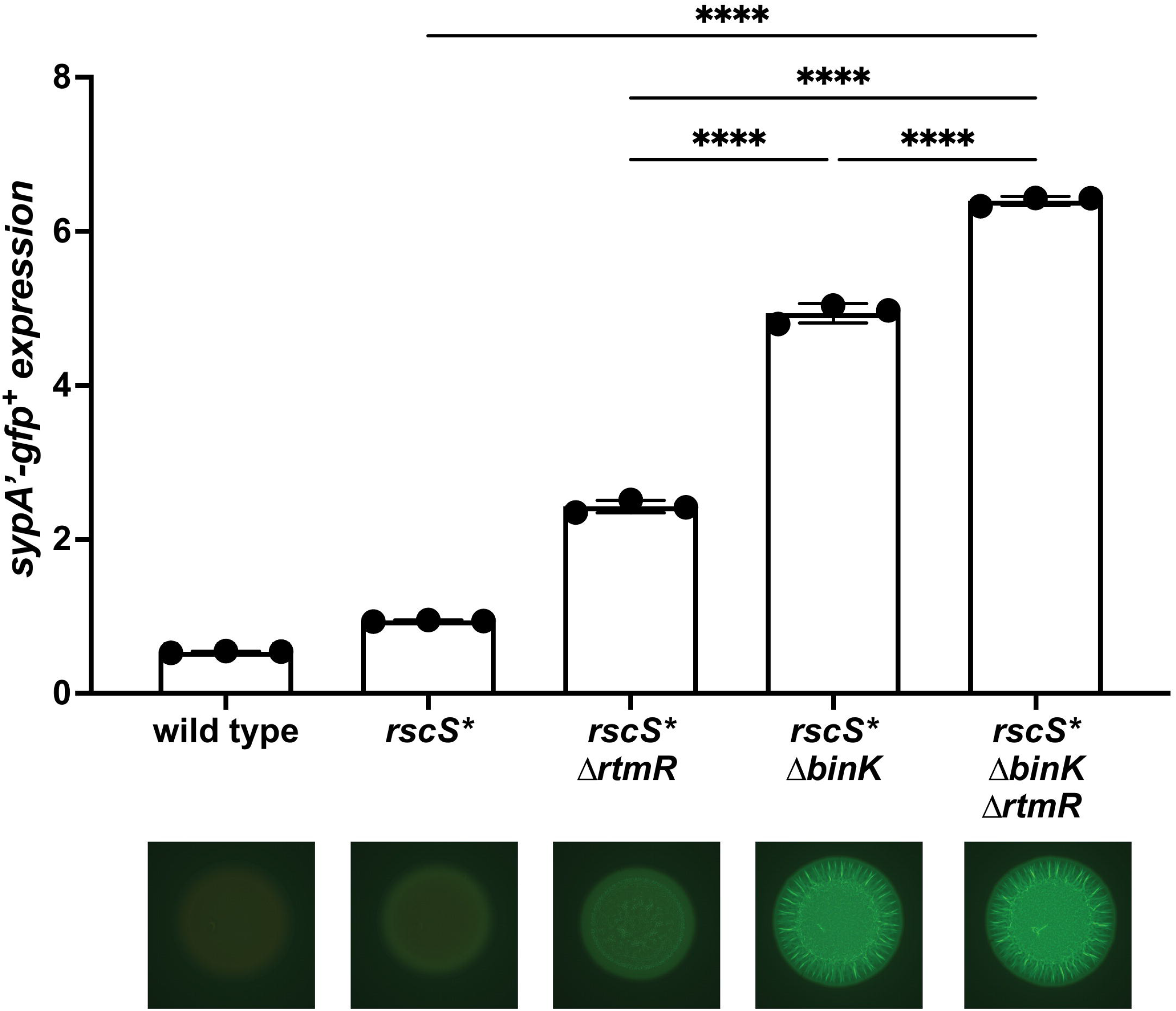
Deletion of *rtmR* and *binK* results in an additive biofilm stimulatory effect. Expression data for the *sypA’-gfp*^+^ reporter in the *rscS\** biofilm model at 28 °C, with single mutations in *binK*, *rtmR*, and a double mutant. Shown are the results for n = 3 biological replicates, collected and analyzed as in Figure 1. ****, P ≤ 0.0001.

### RtmR is a predicted RNA-binding protein with orthologs across the *Vibrionaceae* and related *Gammaproteobacteria*

The chromosomal locus for *rtmR* does not include any known SYP regulators (**Fig. 3A**). Functional annotation of *rtmR* predicted a 160 amino acid protein product with two N-terminal transmembrane helices and a C-terminal RNA Recognition Motif (RRM) domain (**Fig. 3B**). RRM domain proteins have been extensively studied in eukaryotic systems, with functional roles in multiple aspects of RNA processing, including but not limited to tRNA maturation, RNA degradation, and splicing (25–29). Less is known regarding their regulatory roles in bacteria despite their prevalence (30–32).

**Figure 3.**
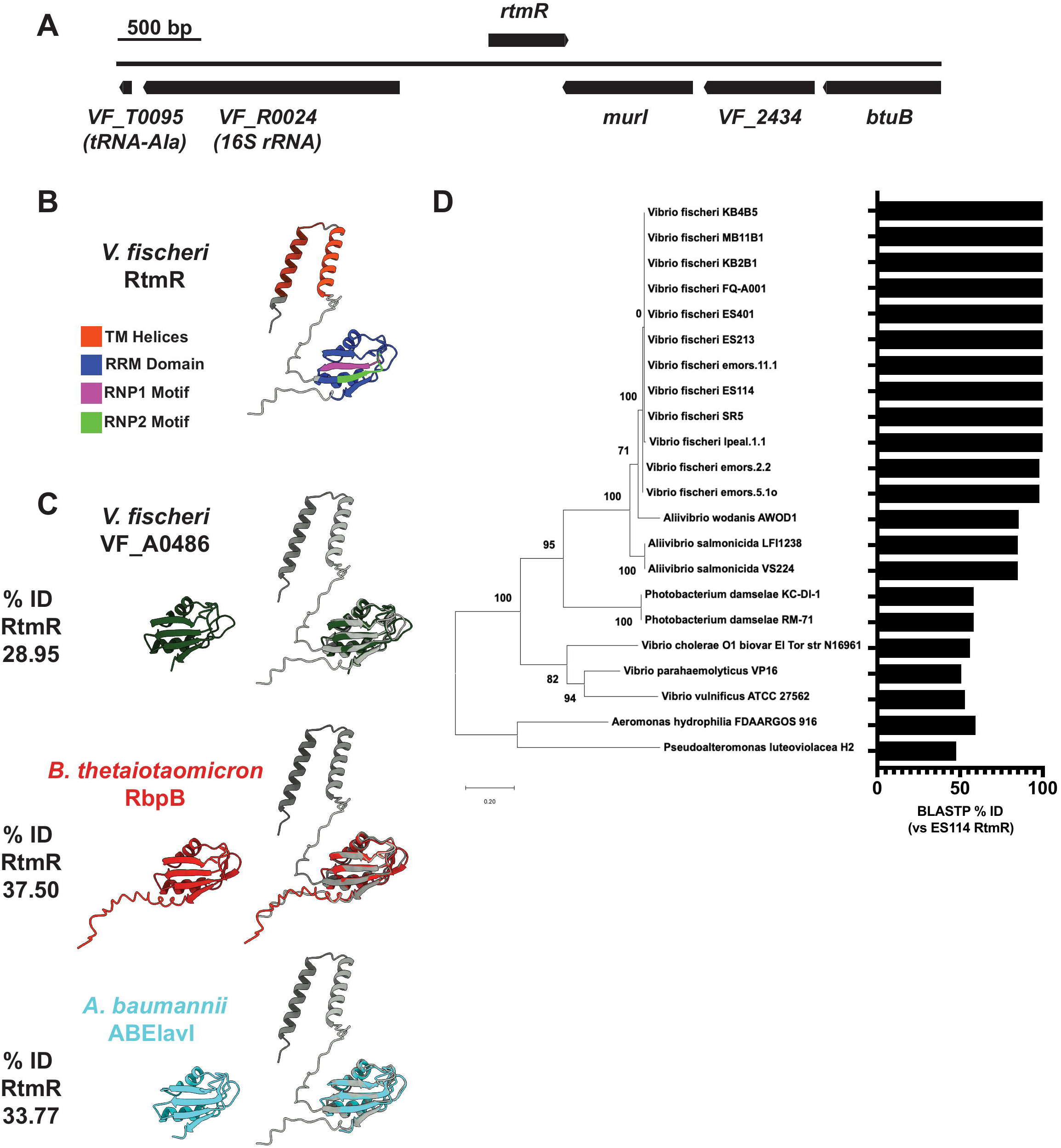
RtmR is a predicted RRM domain RNA-binding protein. A, Locus diagram of *rtmR* and flanking genes, drawn to scale. Genes encoded on the positive strand are on the top of the diagram, and genes encoded on the negative strand are displayed below. B, The corrected RtmR amino acid sequence was submitted to AlphaFold2 ColabFold v1.5.5, and labeled as per domain prediction from InterProScan. Structural predictions were visualized with UCSF ChimeraX. C, Other bacterial RRM domain proteins were modeled with AlphaFold2 ColabFold v1.5.5, and aligned to RtmR (gray) using the UCSF ChimeraX Matchmaker tool. None were predicted to encode transmembrane helices. NCBI BLASTP was used to determine the percent ID of each RBP (query) to RtmR (subject). D, RtmR-specific homologs were identified with TBLASTN and validated for the presence of transmembrane helices using InterPro. For these hits, MEGA11 was used to build a protein phylogenetic tree of the predicted RtmR homologs. Bootstrap support frequencies are shown at each node. Percent ID to RtmR is displayed for each homolog as a bar graph, determined by BLASTP.

There is a second RRM domain protein predicted in *V. fischeri* ES114, VF_A0486, although it lacks any predicted transmembrane regions. Similarly, RbpB in *Bacteroides thetaiotaomicron* and ABElavI in *Acinetobacter baumannii* (30, 33) are both bacterial RRM domain proteins with confirmed RNA-binding activity, and they also lack transmembrane helices (**Fig. 3C**). We conducted TBLASTN searches to identify potential homologs of RtmR beyond *V. fischeri* ES114, yielding >500 hits. Of these, we selected a subset of 21 hits distributed throughout the *Gammaproteobacteria,* all of which had predicted transmembrane helices and an RRM domain, to generate a protein phylogenetic tree of RtmR homologs representing 11 squid-colonizing strains of *V. fischeri* and 10 other species (**Fig. 3D**). Homologs of RtmR appeared to be highly conserved among *V. fischeri* strains: e.g., 8 out of 11 squid-colonizing isolates have a 100% identical RtmR protein (**Fig. 3D**). In all homologs shown in Figure 3D, even where the amino acid identity dropped < 60%, the RtmR homolog was encoded convergently with a gene encoding a glutamate racemase, encoded as *murI* in *V. fischeri*, with broad conservation across the locus.

### RtmR localizes to the membrane

While previous literature has indicated that bacteria do have the capability to localize specific RBPs and RNAs to the inner membrane (34, 35), the presence of transmembrane helices on RtmR was unusual for an RRM domain protein. To assess the membrane localization of RtmR, we conducted cellular fractionation of *V. fischeri* essentially as described previously (36). Signal for RtmR was detectable in total lysate and the membrane fraction of *V. fischeri*, yet no signal for RtmR-3XFLAG was observed in the cytoplasmic fraction, supporting the localization of RtmR as a membrane-bound protein (**Fig. 4**). As a control we assayed for localization of cytoplasmic RNA polymerase α subunit (RpoA). We detected RpoA predominantly in the total lysate and the cytoplasmic fraction. A weak RpoA signal was detectable in the membrane fraction, suggesting that there was some contamination of the cytoplasmic contents into the membrane fraction in our preparations. However, detection of RpoA, but not RtmR, in the cytoplasm provides strong support that RtmR is indeed localized to the inner membrane.

**Figure 4.**
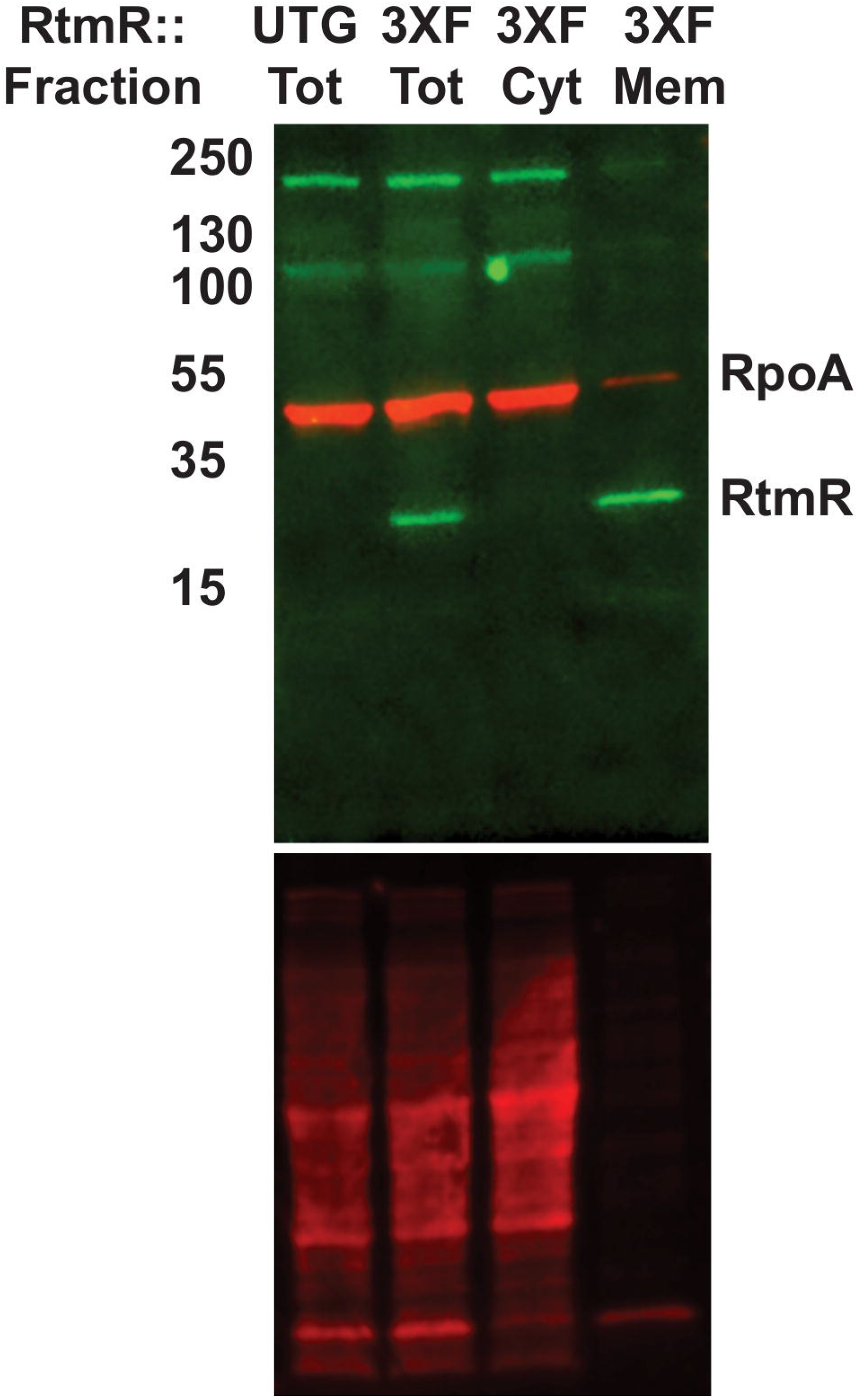
RtmR is detectable in the membrane fraction of *V. fischeri*. Total lysates (Tot) of *V. fischeri* were prepared from strains expressing untagged (UTG) or 3XFLAG-tagged (3XF) alleles of RtmR from single copy complementation constructs inserted in the V. fischeri chromosome at the *att*Tn*7* site. The 3XF strain was fractionated using the method of Matsuda *et al*. (2019) into cytoplasmic (Cyt) and membrane (Mem) fractions following ultracentrifugation at 100,000 x g. Total lysates and cellular fractions were run in a Mini-Protean 4-20% TGX SDS-PAGE gel, and transferred to a 0.45 µm PVDF membrane. Membranes were stained for total protein (bottom panel) using Revert 700 total protein stain, and immunoblotted for the presence of RtmR or RpoA, with RpoA expected to be cytoplasmically localized. Blots were imaged on a LI-COR Odyssey FC imager, and the presented blot is representative of two biological replicates. Band size is indicated in kD.

### The transmembrane helices of RtmR are required for stability of the protein

After detecting enrichment of RtmR in the membrane fraction of *V. fischeri*, we next examined whether membrane localization influences the function of RtmR. We generated two truncation mutants of RtmR lacking the transmembrane helices (**Fig. S2A**). The first truncation removed residues 2-52 (ΔTM-53), eliminating the transmembrane helices yet leaving the linker domain intact. The second mutant truncated RtmR further to leave intact only the RRM domain itself and its C-terminal tail (ΔTM-72), mimicking the structure of VF_A0486 and other known bacterial RRM domain proteins predicted to be localized in the cytoplasm (**Fig. 3B,C**). To assess the stability of the truncation mutants, we compared their expression to that of a wild-type allele driven by the same promoter using immunoblotting (**Fig. S2B**). While the lysates of the truncation mutants were loaded similar to the wild type as assessed by total protein staining, we were unable to detect any RtmR signal from the truncation mutants. As would be expected based on the immunoblotting results, both membrane truncation mutants exhibited weak if any biofilm inhibition at 28 °C (**Fig. S2C**). These results suggest that the transmembrane helices of RtmR, or potentially the transcript encoding them, play a critical role in the stability or synthesis of the protein.

### Mutagenesis of the RRM domain disrupts RtmR biofilm regulation

RRM domains maintain a conserved fold, forming a single β-sheet with four antiparallel β-strands backed by two α-helices (β_1_α_1_β_2_β_3_α_2_β_4_) (37). RtmR’s RRM domain maintains both motifs with strong conservation, including all amino acids known to be involved in direct RNA contact (**Fig. 5A**) (38). We tested whether disruption of these conserved motifs impacted RtmR activity through generating an RtmR^RNPnull^ allele by introducing substitutions in RNP1 (R112Q, F114V, F116V) and RNP2 (Y74V), all established mutations for deactivating the RRM domain (39, 40). We introduced *rtmR^RNPnull^* into the chromosome of *rscS\** Δ*rtmR* at the neutral site used for all complementations (*att*Tn*7*). Compared to the parent unmutated allele of RtmR, RtmR^RNPnull^ permitted biofilm formation earlier at 28 °C in wrinkled colony biofilm assays, reflected by increased colony wrinkling by 24 hrs (**Fig. 5B**). The RtmR^RNPnull^ strain had substantially increased *syp* expression (**Fig. 5C**). Both wild-type RtmR and RtmR^RNPnull^ alleles were equally expressed, suggesting that the observed loss of activity in RtmR^RNPnull^ was unlikely to be due to decreased expression or stability (**Fig. S3**). These results support the hypothesis that the conserved RNP1 and RNP2 RNA-interaction residues are required for RtmR functionality and provide support that RtmR acts as an RNA-binding protein.

**Figure 5.**
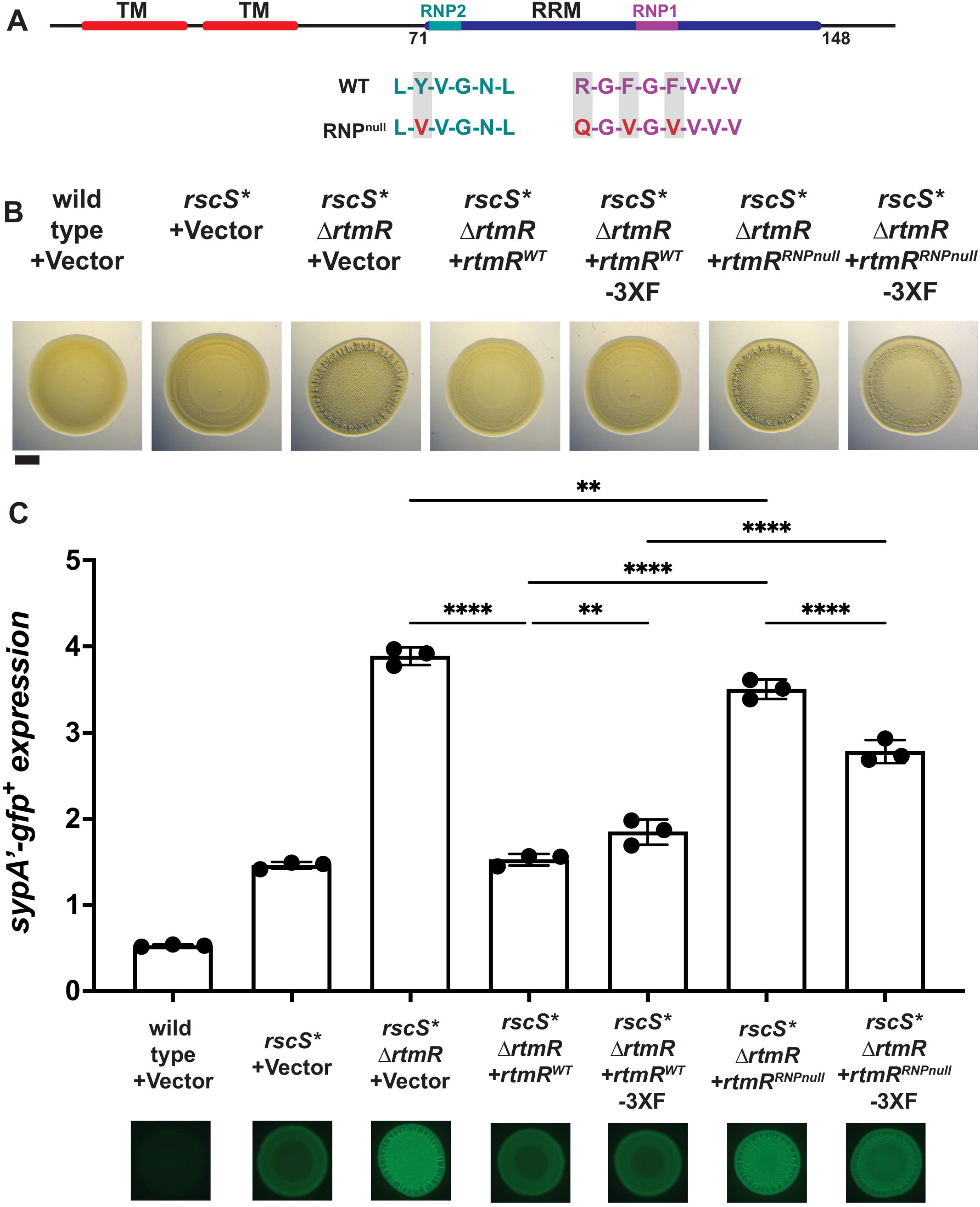
RtmR requires intact RRM motifs for biofilm inhibition. A, Predicted RtmR RNP1 and RNP2 motifs are provided below a scaled diagram of the RRM domain. Start and end residues of the domain are provided. Residues expected to act in RNA-binding are shaded in gray. Amino acid substitutions used to generate the RNP-dead (RNPnull) mutant are provided below the wild-type sequence in red. B, Cultures of indicated strains were spotted on LBS agar and allowed to grow at 28 °C. At 24 h colony morphology was recorded using a Leica M60 microscope. The + symbol indicates single-copy complementation at the chromosomal *att*Tn*7* site with the vector control or *rtmR* allele. Scale bar represents 5 mm. C, Colony spots carrying a plasmid-based *sypA’-gfp*^+^ transcriptional reporter were assayed for *syp* locus expression at 28 °C after 24 hrs of growth. Representative colony fluorescence is shown for each strain, and the assay was conducted with n = 3 biological replicates, each shown here as the average of two technical duplicates. Statistical analysis of *syp* expression was conducted in Prism 10, using an ordinary one-way ANOVA with Šidák’s multiple comparison test. ****, P ≤ .0001, **, P ≤ .01.

### RtmR is an RNA-binding protein

Given the importance of RtmR’s RNA-binding residues, we sought to directly validate RNA-binding activity and identify any bound RNA ligands. An individual-nucleotide resolution crosslinking and immunoprecipitation sequencing (iiCLIP-seq) approach (41) was adapted to *V. fischeri*. Following growth to early stationary phase (OD600 ∼1.8) with the functional 3XFLAG-tagged RtmR allele (**Fig. 1**), UV-irradiation was used to form covalent crosslinks between bound RNAs and RtmR, after which bound RNAs were trimmed by partial RNase digestion to enable sequencing of the bound RNAs. RtmR-RNA complexes were then immunoprecipitated and stringently washed to remove unbound RNAs. Co-precipitated RNAs were 3’-labeled with a fluorescently-tagged DNA adapter to allow visualization and generation of sequencing libraries. Immunoprecipitations resolved a labeled RtmR-RNA complex at the predicted size of monomeric RtmR-3XFLAG + DNA adapter (∼35 kD), with two further complexes also detected at higher molecular weights (**Fig. 6**). These larger complexes were of equivalent size to RtmR complexes identified in a preliminary immunoprecipitation assay, suggesting that all of the bands identified represented RNA-bound RtmR (**Fig. S4**). These complexes would only have been observed if RtmR was able to bind RNA, as the fluorophore was specifically ligated to the 3’ end of bound RNAs, validating that RtmR does engage RNA ligands inside live cells. RNA:protein complexes were absent in controls demonstrating the specificity of our approach. We recovered the bands shown in **Figure 6** for Illumina sequencing and identification of the bound ligands.

**Figure 6.**
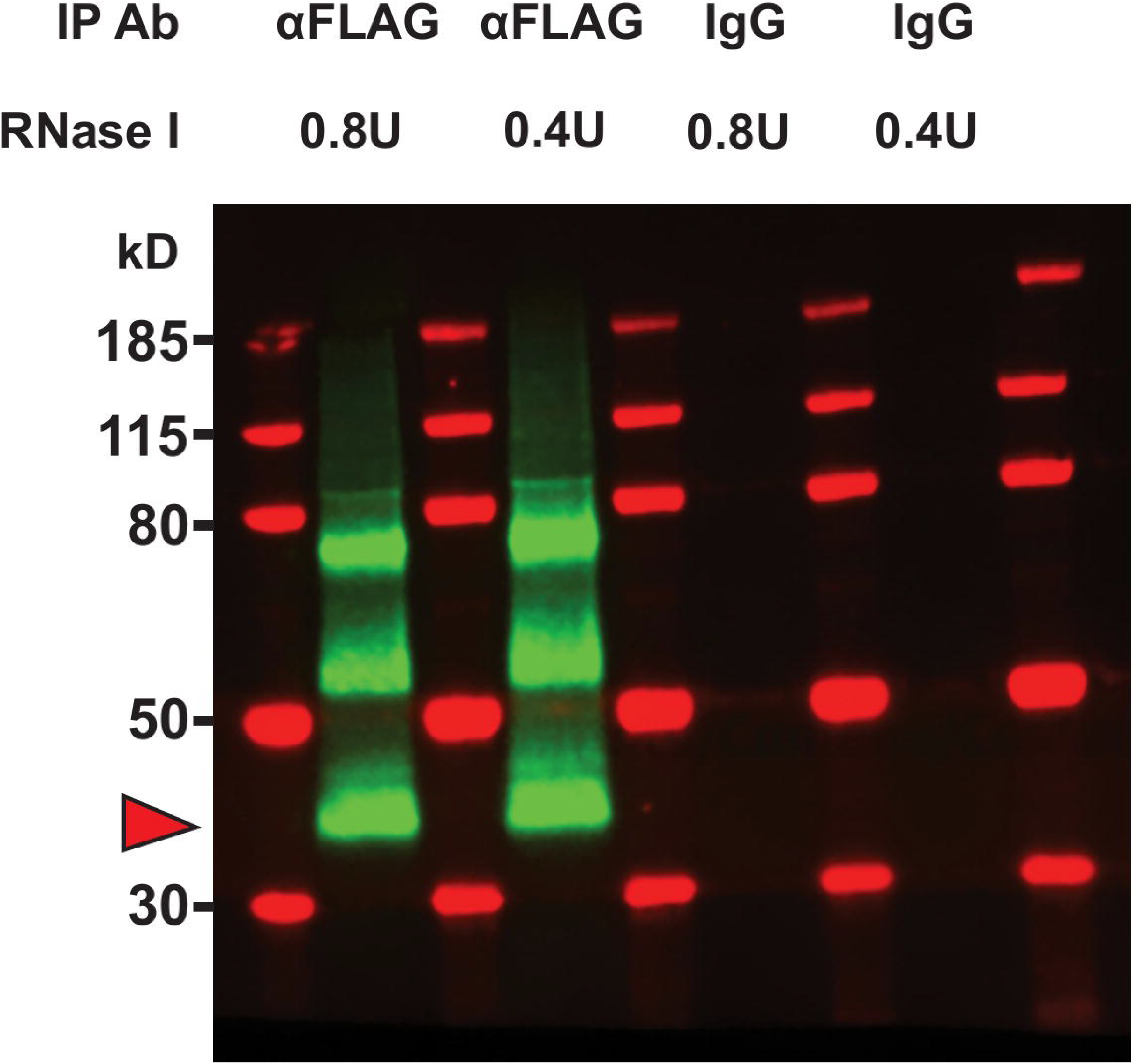
RtmR binds RNA as part of multiple complexes. A, *V. fischeri* strain carrying a RtmR-3XFLAG allele expressed from a single-copy complementation construct inserted at the *att*Tn*7* site was grown to early stationary phase, followed by exposure to 800 mJ/cm2 UV irradiation. Cells were lysed and immunoprecipitated with either a monoclonal mouse anti-FLAG antibody or non-specific mouse IgG. RNA ligands bound by immunoprecipitated RtmR were 3’ labeled with the L3-IR-App fluorescent DNA adapter. RNA-RtmR complexes were then run in a 4-12% NuPAGE Bis-Tris gel, and immunoblotted onto 0.45 µm Protran nitrocellulose membranes, followed by imaging with a LICOR Odyssey FC imager. Shown here is representative of n = 2 biological replicates. Red arrowhead indicates the predicted size of monomeric RtmR carrying an RNA labeled with the L3-IR-App adapter.

### RtmR has a broad RNA interactome

The output from the RtmR iiCLIP-seq experiment was analyzed using the nf-core/clipseq pipeline, enabling the discovery of RtmR-ligand interaction sites at single nucleotide resolution based on protein-RNA crosslink positions (42). Regions of significant crosslink enrichment (peaks) were determined by the software tool iCount, after which peak scores were then summed on a per gene basis to assess transcript enrichment in the iiCLIP-seq output. This analysis captured a total of 1588 transcripts interacting with RtmR across both replicates (**Table S1**), of which 383 transcripts had greater than 100 peak score in both replicates. RtmR-interacting ligands were quite diverse including mRNAs, a majority of the known *V. fischeri* small RNAs, and tRNA/rRNA transcripts (**Fig. 7A**).

**Figure 7.**
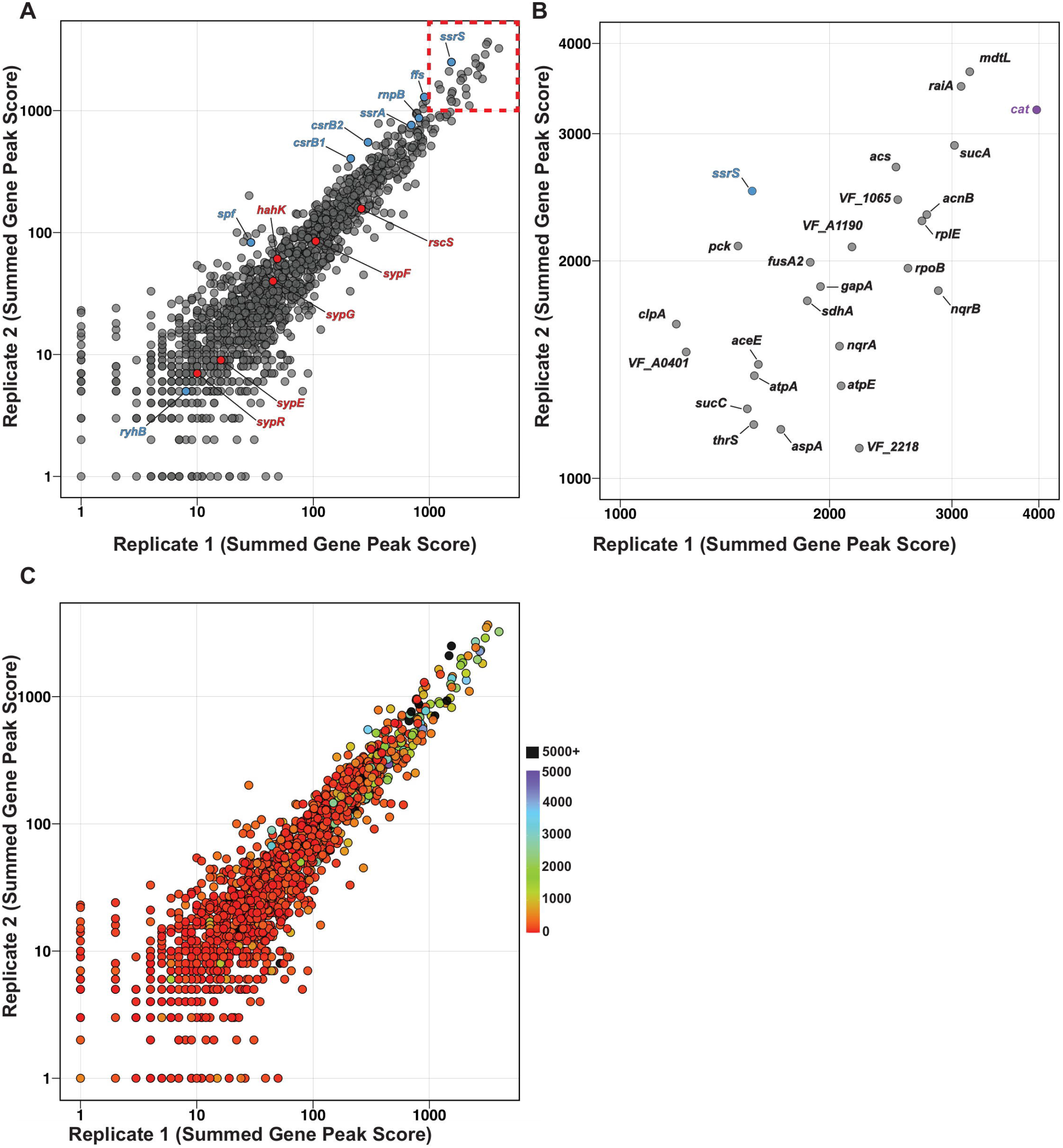
RtmR is a promiscuous RNA-binding protein. A, To profile ligand enrichment across both replicates in the iCLIP-Seq experiment, crosslink counts from statistically significant peaks predicted by iCount were summed within known *V. fischeri* transcripts (peak score). For ease of viewing the data, the X- and Y-axes are log-scaled. Transcripts from small RNA genes are labeled in light blue, and genes belonging to SYP regulation are colored in red. Gene annotations for labeled genes are provided. The area with the most abundant shared ligands is indicated with a dashed box. B, The most abundant ligands detected by the iCLIP-seq, as shown in A’s dashed box, with labels. C, Ligands were plotted according to crosslink abundance as in A, with gradient coloration of ligand expression as measured by Transcripts per Kilobase Million (TPM) in the RNA-sequencing assay described in Fig. S5.

After observing the broad binding profile of RtmR, we hypothesized that RtmR could have wide-ranging effects on transcripts in *V. fischeri*, and/or that the distribution could be explained by non-specific binding with the most enriched ligands matching the transcripts with highest abundance in the transcriptome. Therefore, we conducted RNA-seq analysis under the same conditions as the iiCLIP-seq (early stationary phase; OD600 ∼1.8) and examined the effect of RtmR on the bacterial transcriptome. Comparing a Δ*rtmR* mutant against an isogenic wild-type ES114 strain (**Fig. S5AB**) revealed differential expression of greater than 400 genes, under thresholds of | ≥ 1 | Log_2_(Fold Change) and False Discovery Rate (FDR) < 0.05 (**Table S2**). We integrated the data from the bulk RNA-seq, measuring gene expression through TPM, for each ligand in the iiCLIP-seq dataset (**Fig. 7C and Table S1, S4**). The top iiCLIP ligands were not the most expressed transcripts in the cell, suggesting some means of specificity to RtmR activity. However, attempts to discern a binding motif using DREME and STREME failed to yield high confidence predictions. Without a reliable motif from these programs, we turned to directly visualizing iiCLIP binding patterns across the top 8 ligands in the case that specific regions of engagement could be mined for cryptic motifs. We detected well-defined binding peaks of RtmR in some instances (*sucA, mdtL*, *raiA*), while others had a pattern that suggested that RtmR was broadly engaged across the length of the transcript (e.g., *VF_1065,* and *acnB*) (**Fig. S6**).

### RtmR biofilm inhibitory activity requires overexpression of the sensor kinase RscS

To understand RtmR’s biofilm-inhibitory phenotype, we examined the iiCLIP-seq results to determine whether known SYP regulators were bound. Biofilm biosynthetic gene *sypR* and biofilm regulator genes *sypE, sypF, sypG, binK, hahK*, *and rscS* were present in the dataset. Most were low-abundance hits with less than 100 peak score, with the exception of the biofilm activator *rscS* (**Fig. 7A**). The strain used in the iiCLIP-seq assay carried the *rscS\** allele, wherein a Tn*5* transposon is integrated 34 bp upstream of the *rscS* open-reading frame. Increased expression of *rscS* in this model is driven from the promoter of the FRT site-flanked chloramphenicol acetyltransferase (*cat*) gene, originally derived from pKD3, on the transposon (22).

Media supplementation of seawater relevant levels of calcium (10 mM) is sufficient for sensor kinase HahK to aid *syp* induction through SypF in a largely RscS-independent fashion, and has previously been used as a model of biofilm formation in *V. fischeri* (**Fig. 8A**) (43–45). If the biofilm regulatory target of RtmR was RscS-independent, we expected to observe increased biofilm formation in the calcium model similar to what we had observed in *rscS\**. As previous literature had identified that induction of biofilm in the calcium model was achievable in the Δ*binK* background (44), we used a Δ*binK* mutant as a positive control. Further, given that we had observed a partial additive effect of deleting both *binK* and *rtmR* in the *rscS\** model, we assayed a double mutant in the calcium model. Only strains carrying the Δ*binK* allele (either alone or in the double mutant) were capable of forming wrinkled colony biofilms in the calcium model, with no effect observed of Δ*rtmR* in either the wild type or Δ*binK* background (**Fig. 8B**). Measurement of *syp* transcription within colony spot biofilms revealed that Δ*rtmR* did not induce *syp* expression compared to wild type in the calcium model, while the Δ*binK* mutant was competent to induce substantially higher *syp* expression as expected (**Fig. 8C**). In the Δ*binK* background, removal of RtmR led to approximately 20% reduced *syp* expression, demonstrating an unexpected role for RtmR in promoting biofilm transcription under these conditions (**Fig. 8C**). However, it was clear that the major phenotype we observed for RtmR in inhibiting biofilm formation occurred only in the *rscS\** model that was RscS-dependent.

**Figure 8.**
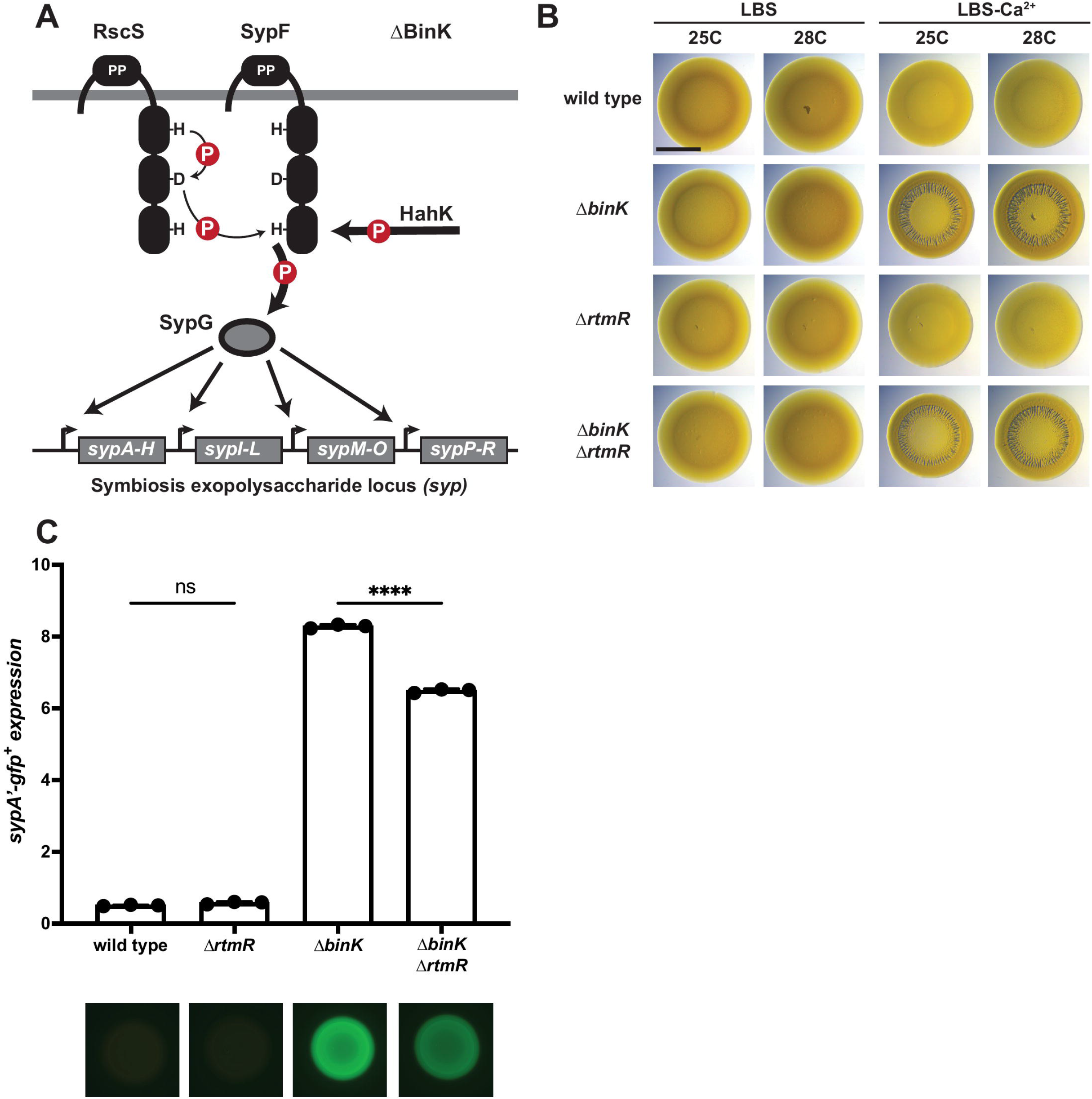
RtmR biofilm regulation is specific to the RscS overexpression model. A, Regulatory diagram of the Δ*binK*-Ca^2+^ biofilm model, predicted contributions of phospho-transfer networks indicated through arrow thickness. B, *V. fischeri* strains were spotted on nutrient agar with or without 10 mM calcium supplementation, and allowed to grow at 25 or 28 °C. After 48 hrs, colonies were visualized on a Leica M60 microscope. C, Assessment of syp locus expression within colony spots grown on calcium-supplemented nutrient agar was conducted with a plasmid-based *sypA’-gfp*^+^ transcriptional reporter as described previously. Shown are the results of n = 3 biological replicates, each the average of two technical replicates. Comparisons of *syp* expression were conducted in Prism 10 using an ordinary one-way ANOVA with Šídák’s multiple comparisons test. Representative colony images are provided for each strain. ns, not significant. ****, P ≤ 0.0001.

### RtmR does not regulate biofilm formation via the heterologous *cat* gene

In the *rscS\** model, expression of *rscS* is likely driven from the *cat* promoter. In the iiCLIP-seq we detected the heterologous *cat* gene as one of the most enriched RtmR ligands (**Fig. 7A, S7**). Given the key role of *rscS* as the primary activator of biofilm formation in the *rscS\** model, as well as the enrichment of *rscS* over other biofilm regulators and the exceptionally high number of *cat* interactions in the iiCLIP-seq, we tested whether the RtmR biofilm-inhibitory phenotype was linked to the *cat* gene. In the *rscS\** genetic background we generated a Δ*cat* allele (**Fig. S8A**), and we assessed the ability of the mutant to form biofilms at 25 °C and 28 °C in the presence and absence of *rtmR* (**Fig. S8B**). In the Δ*cat* background, RtmR contributed a similar biofilm inhibition phenotype as the parent strain, and Δ*cat* Δ*rtmR* showed clearly improved biofilm formation at 28 °C compared to 25 °C. We did note that the *rscS\** Δ*cat* background had overall lower levels of biofilm than the *rscS\** parent, likely due to baseline effects on *rscS* transcription.

Therefore, the biofilm inhibitory effect of RtmR is not *cat* dependent. While we did not detect a role for *cat* in biofilm regulation, the high enrichment of this transcript in the iiCLIP-seq assay raised the possibility that RtmR may also regulate chloramphenicol resistance in strains encoding *cat.* Using a plate-based growth assay, we determined that deletion of *rtmR* resulted in reduced levels of chloramphenicol resistance in the *rscS\** model (**Fig. S9**).

### RtmR constrains RscS expression

The comparison of the *rscS\** and calcium models had suggested that RtmR biofilm regulation was RscS-dependent, and iiCLIP-seq revealed RtmR binding at the 5’ end of the *rscS* CDS and within co-transcribed *cat*. As described above, RtmR-*cat* interactions did not explain the biofilm regulatory phenotype of RtmR. Following this result, we refocused to studying interactions of RtmR with *rscS* itself. RNA-seq data comparing *rscS\** Δ*rtmR* to *rscS\** Δ*rtmR rtmR-*3XFLAG did not detect differential expression of *rscS* (**Fig. S5CD, Table S3**). Given the known capability of other bacterial RBPs to regulate the translation of mRNA ligands (46–48), we examined whether RtmR could influence levels of RscS using immunoblotting (**Fig 9**). Comparing RscS levels in *rscS\** and *rscS\** Δ*rtmR* revealed an increase in RscS expression in the Δ*rtmR* mutant (**Fig. 9A Lane 3 and 4**). As done for previous Δ*rtmR* mutant strains, we inserted *rtmR* in single copy at the *V. fischeri* chromosomal *att*Tn*7* site to evaluate this phenotype upon complementation (**Fig. 9A Lane 8**). We also generated a strain carrying an intact native *rtmR* locus and an additional copy of *rtmR* at the *att*Tn*7* site in the *rscS\** genetic background. This second strain served to assess whether increasing the gene dosage of *rtmR* would result in a further drop in RscS-3XFLAG levels (**Fig. 9A Lane 7**). Compared to strains carrying empty insertions at *att*Tn*7* (**Fig. 9A Lane 5 and 6**), we noted that complementation with the single-copy allele reduced RscS levels to that of the parent strain, while increasing the gene dosage of *rtmR* did not further reduce RscS levels (**Fig. 9AB**). These results revealed that RtmR is capable of reducing the expression of biofilm regulator RscS.

**Figure 9.**
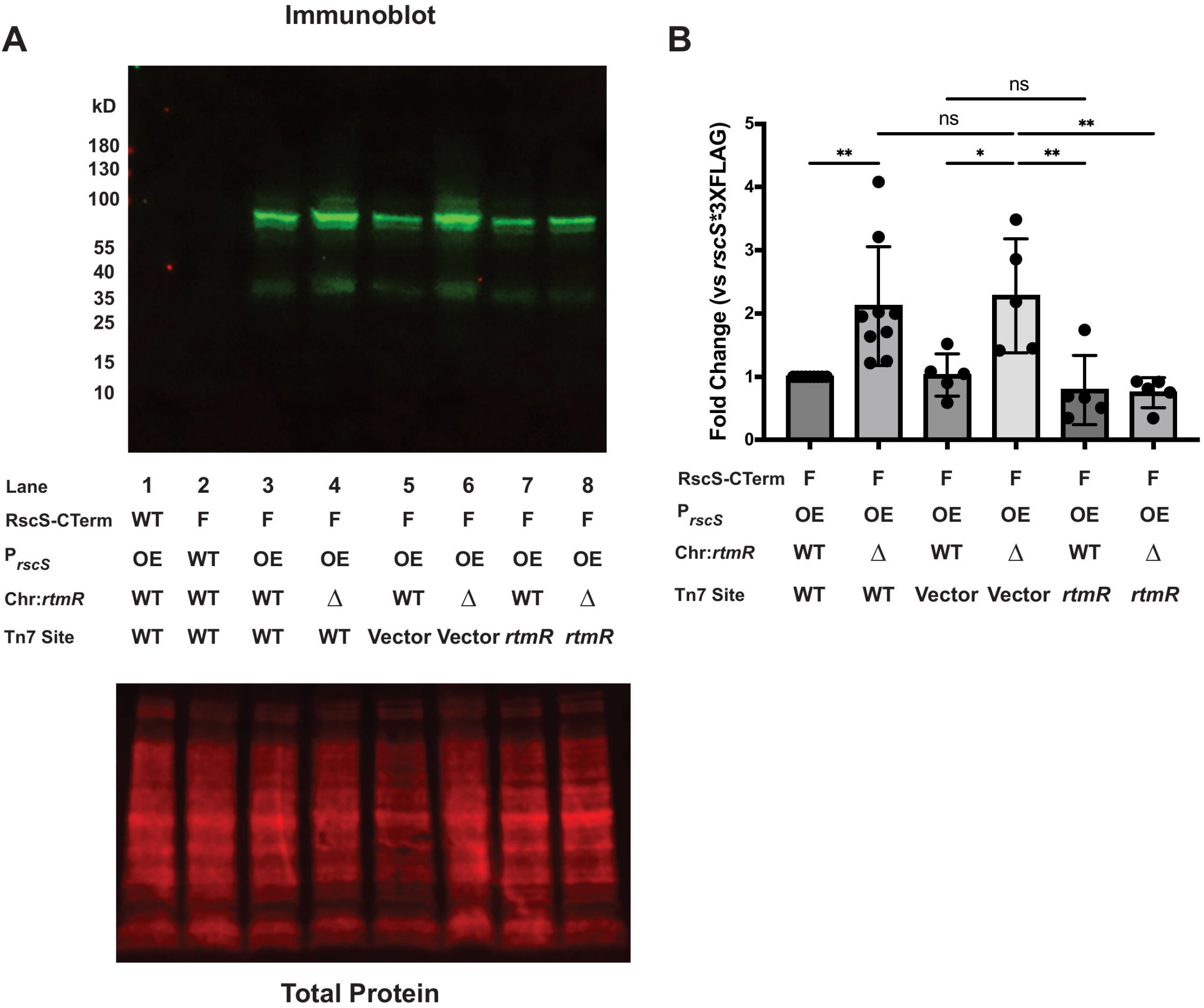
RtmR reduces the expression of the sensor kinase RscS. A, *V. fischeri* strains with or without a 3’ 3XFLAG tag appended to *rscS* were grown overnight at 28 °C. The following morning, cultures were lysed in 1% SDS with protease inhibitors and the total protein yield of each lysate was measured. Lysates were then normalized for gel loading and run through a 4-20% Tris-Glycine SDS-PAGE gel, followed by wet transfer to a 0.45 µm PVDF membrane. Efficiency of transfer and overall loading was assayed using the LI-COR Revert 700 total protein stain, followed by immunostaining with a monoclonal mouse anti-FLAG antibody and a goat anti-mouse IgG IRDye 800CW secondary antibody. Shown is a representative blot (top panel) with respective total protein stain (bottom panel). RscS-CTerm is the status of the RscS C-terminus either wild type (WT) or 3XFLAG-tagged (F). P*rscS* is the promoter status of *rscS*, either wild type (WT) or the *rscS*\* overexpression allele (OE). Chr:*rtmR* indicates whether *rtmR* is present at the native (WT) wild type locus or deleted (Δ). *att*Tn*7* site is a neutral site in the first *V. fischeri* chromosome at which the *Tn7* transposon specifically inserts, there may be either no *Tn7* transposon insertion (WT), insertion of a vector control empty *Tn7* transposon (Vector), or a *Tn7* transposon carrying a complementation allele of *rtmR* (*rtmR*). B, Expression levels of RscS-3XFLAG were measured from blots as shown in A, with RscS signal (∼100 kd band) normalized to total protein staining. Measurements were taken in LICOR ImageStudio, and statistical analysis was conducted in Prism 10, using an ordinary one-way ANOVA with Šídák’s multiple comparisons test. ns, not significant. *, P ≤ 0.05. **, P ≤ 0.01.

## DISCUSSION

RNA-based regulation is well established for many members of the *Vibrionaceae*, exemplified by the central role of Quorum regulatory RNA (Qrr) sRNAs in mediating the transition between low and high cell density in the *Vibrio* quorum sensing system (49–53). The squid symbiont *V. fischeri* is rich in known RBPs including Hfq (VF_2323), CsrA (VF_0538), ProQ (VF_1279), and multiple Cold shock proteins (VF_A0595 and VF_2561). Several of these proteins have been demonstrated to regulate phenotypes relevant to host colonization, including regulation of bioluminescence by Hfq and CsrA (54–57), flagellar motility by Hfq (57, 58), and regulation of multiple exopolysaccharide synthesis pathways by Hfq (57). *V. fischeri* also possesses predicted RBPs that had not been examined, such as VF_2432 and VF_A0486. In this work, we used an *in vitro* biofilm model to capture activity of one of these regulators, VF_2432 (RtmR). RtmR is representative of a new class of bacterial RBP, directly combining an RRM RNA-binding domain with integrated transmembrane helices. We demonstrate that RtmR requires conserved RNA-binding motifs in the RRM domain for full functionality, and observe RNA binding on immunoprecipitated RtmR. To further emphasize the importance of the RRM domain to RtmR function, the addition of a 3XFLAG tag proximal to the RRM domain imparted a slight defect in biofilm regulation. RNA ligands isolated from RtmR were diverse, suggesting that RtmR may act as a global post-transcriptional regulator in *V. fischeri.* For one of these ligands, the biofilm sensor kinase RscS, we used immunoblotting to reveal that levels of the product are regulated inversely to levels of RtmR. This interaction with RscS is the basis of our current model of RtmR biofilm regulation, wherein control of RscS levels impacts stimulation of the biofilm *syp* EPS locus (**Fig. 10**). We also detected a strong interaction between RtmR and transcripts from a bioengineered chloramphenicol resistance gene, *cat*. In a strain carrying *cat*, deletion of *rtmR* resulted in a growth defect on chloramphenicol-supplemented media. Future studies will be necessary to determine if RtmR plays a role in regulating this and other antibiotic resistance genes, or horizontally acquired genes more broadly.

**Figure 10.**
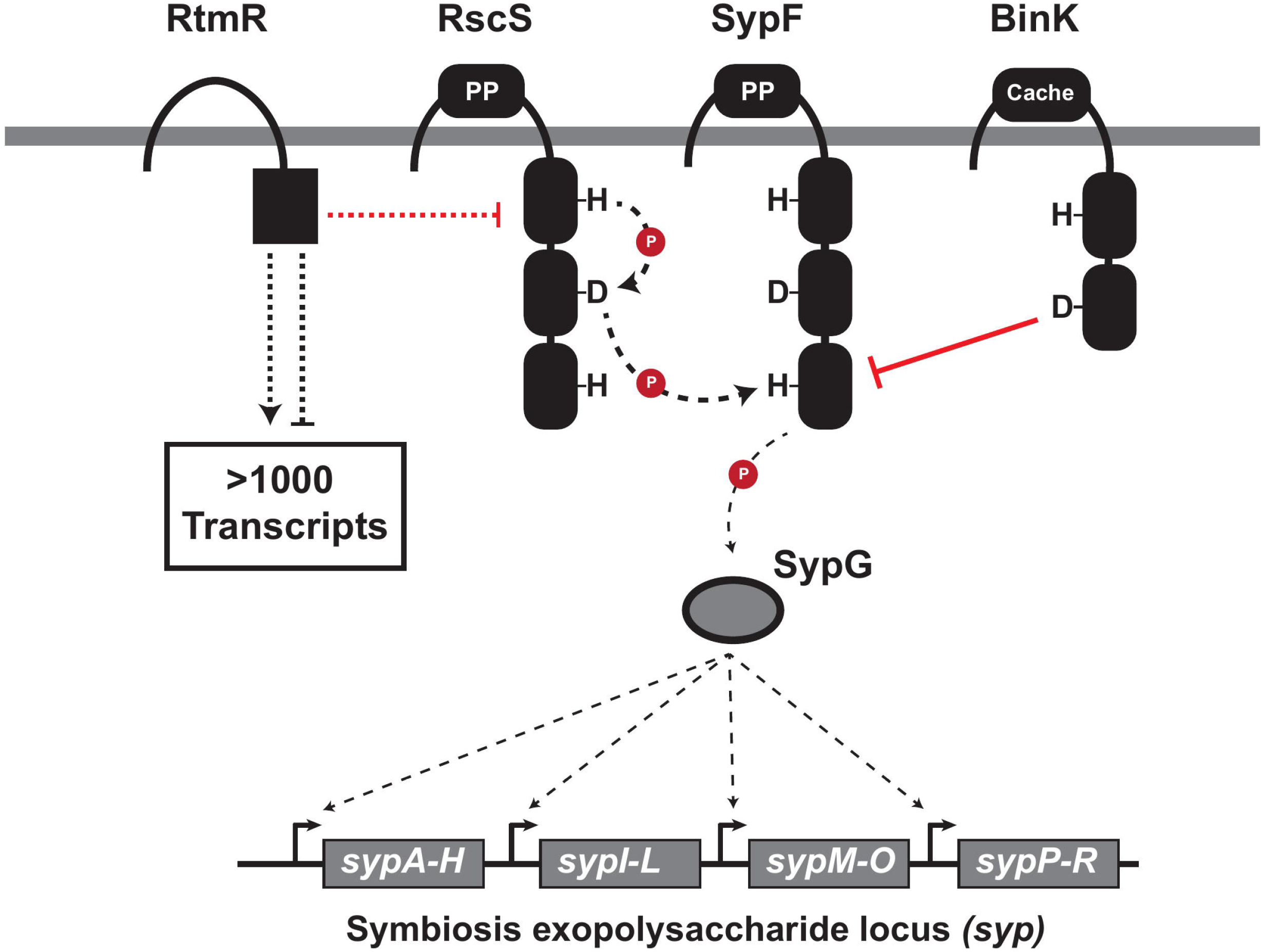
RtmR is a promiscuous RNA-binding protein with integration into Syp regulation. Presented above is the current working model of biofilm regulation by RtmR. RtmR, via an unknown mechanism, functions as a negative regulator of RscS levels. Reduced levels of RscS result in lower stimulation of the Syp phosphorelay, compounding upon the inhibitory effects of the other known inhibitor of the Syp phosphorelay, BinK. The downstream response regulator SypG therefore remains predominantly unphosphorylated, resulting in far weaker activation of the biofilm syp exopolysaccharide locus’ expression. Alongside this effect, RmtR interacts with more than 1000 transcripts as determined by CLIP-seq suggesting further regulatory effects which remain to be discovered. Abbreviations and symbols: PP, Periplasmic domain, Red “P”, phosphoryl group.

RRM domain proteins are abundant in all three domains of life, with extensive characterization in eukaryotic systems (26, 27, 40, 59–64). However, few bacterial RRM domain proteins have received attention compared to the literature on other bacterial RBPs such as Hfq, ProQ, and CsrA, leaving their functional roles mysterious. The RbpB RRM domain protein of *Bacteroides thetaiotaomicron* is a regulator of polysaccharide metabolism (**Fig. 3C**), and is composed of an RRM domain with an unstructured C-terminal tail (30, 32). RbpB binds single stranded RNA (ssRNA) with some degree of specificity, as measured by electrophoretic mobility shift assays (EMSAs) of RbpB in complex with ssRNA probes carrying all possible pentamer sequences (30). Follow up studies with RbpB has confirmed via CLIP-seq analysis that it has broad RNA-binding activity of both mRNAs and sRNAs (32), and the Rbp proteins facilitate growth at cold temperatures (65).

Our data suggest that RtmR is an RBP with promiscuous RNA binding, with iiCLIP-seq yielding >1000 mRNA ligands. Our attempts to identify an RNA-binding motif through DREME (66) and STREME (67) with RtmR iiCLIP-seq data did not yield confident sequence predictions. An important limitation to our results is that amid the large number of iiCLIP-seq ligands presented, it will be important to validate individual hits of interest. Despite lacking a clear binding motif, RtmR does appear to be capable of interacting with transcripts of a broad swathe of cellular processes. This activity is also reflected in *B. thetaiotaomicron* Δ*rbp* mutants that have hundreds of differentially regulated genes in RNA-seq (30), supporting a role for bacterial RRM domain proteins as global regulators.

Additionally, the number of RNA ligands of RtmR may be more expansive than captured in our iiCLIP-seq analysis due to two caveats. First, we required each read to map uniquely in the chromosome. This requirement is of particular relevance to rRNA and tRNA ligands, which have high similarity and typically cannot be uniquely mapped. Attempts to visualize the rRNA and tRNA components of our dataset by permitting multi-mapping induced artificial inflation of hits to these loci, hindering our ability to detect the overall level of RtmR interaction with these ligands accurately. Given the localization of *rtmR* near a large rRNA/tRNA locus, these interactions are a focus for future study. The second caveat is that these results reflect the binding of RtmR explicitly to annotated gene features, as the peak calling software used in our study, iCount, did not analyze the crosslink events occurring within intergenic regions. Approximately 14.5% and 16.5% of the total crosslinks in the iiCLIP-seq experiment were localized in intergenic regions in replicates 1 and 2, respectively, suggesting sRNAs that may be useful to pursue in future work.

A feature that RtmR does not share with RbpB or previously characterized bacterial RRM domain proteins are its two N-terminal transmembrane helices, which we were unable to remove without rendering the protein undetectable in the cell. These results may suggest that proper folding of RtmR is dependent on the transmembrane helices, or that the RRM domain is proteolyzed rapidly when not membrane-localized. Alternatively, as we used the putative *rtmR* promoter region to drive both constructs, it is possible that the *rtmR* transcript is post-transcriptionally regulated. The iiCLIP-seq dataset does predict multiple RtmR interaction peaks within the first 100 nt of the CDS, suggesting that RtmR has the potential to auto-regulate through the region encoding the transmembrane helices.

Overall, we report the detection of a novel RNA-binding protein in *V. fischeri*. Our approach leveraged the extensive previous characterization of *V. fischeri* biofilm regulatory pathways to identify essential structural components of RtmR itself and identify possible downstream targets. RtmR is representative of a new class of RBP distributed throughout the *Vibrionaceae*, yet so far these regulators have remained cryptic as to their activities. This work therefore notes the value of the vibrio-squid model as an effective tool in revealing functions of previously unannotated gene products, which may contribute to a better understanding of signal transduction broadly and bacterial RNA regulation specifically.

## MATERIALS AND METHODS

### Bacterial strains, media, and growth conditions

Strains of *Vibrio fischeri* and *Escherichia coli* used in this work are provided in **Table 1**, and plasmids generated for this work are listed in **Table 2**. Unless specified otherwise, *V. fischeri* strains were grown in LB Salt (LBS) media (per liter: 25 g LB Miller powder [BD Difco], 10 g NaCl, and 50 mL 1 M Tris buffer [pH: 7.5]), and *E. coli* strains were grown in LB (per liter: 25 g LB Miller powder [BD Difco]) or brain heart infusion media (BHI) (per liter: 37 g BHI powder [BD BBL]). For solid media, the above growth media were supplemented with 1.5% agar [BD]. Antibiotics were added into growth media as necessary, *V. fischeri*: erythromycin (Erm) (5 µg/mL), kanamycin (Kan) (100 µg/mL), and chloramphenicol (Cam) (2.5 or 5 µg/mL), *E. coli:* erythromycin (150 µg/mL), kanamycin (50 µg/mL), carbenicillin (Carb) (100 µg/mL), and chloramphenicol (25 µg/mL). Conjugation of plasmids into *V. fischeri* was accomplished through standard procedures (68).

**TABLE 1.** Strains used in this work.

| Strain | Genotype | Source or reference |
| --- | --- | --- |
| *Vibrio fischeri* |  |  |
| MJM1100 | Natural isolate, Euprymna scolopes bobtail squid light organ = ES114 | (86, 87) |
| MJM1198 | MJM1100, IG (*glpR-rscS*)::Tn5 (Cam<sup>R</sup>) = *rscS* * | (22) |
| MJM1438 | MJM1100 / pM1422 | (21) |
| MJM1492 | MJM1198 / pM1422 | This work |
| MJM1538 | MJM1100 / pLostfoX | (69) |
| MJM2009 | MJM1198 *rtmR*::Tn<sup>erm</sup> | This work |
| MJM2012 | MJM1198 *binK*::Tn<sup>erm</sup> | This work |
| MJM2251 | MJM1100 $\Delta binK$ | (20) |
| MJM2252 | MJM1100 $\Delta binK$ / pLostfoX-Kan | (20) |
| MJM2479 | MJM1198 *attTn7*::*erm* | (20) |
| MJM2489 | MJM1198 *attTn7*::*erm* / pM1422 | (20) |
| MJM2493 | MJM1100 $\Delta binK$ / pM1422 | (21) |
| MJM3381 | MJM1100 *attTn7*::*erm* | (45) |
| MJM4627 | MJM1100 $\Delta rtmR$::*erm-bar* | This work |
| MJM4639 | MJM1100 $\Delta rtmR::bar$ | This work |
| MJM4674 | MJM1100 $\Delta rtmR::bar attTn7::P_{rtmR}-rtmR-erm$ | This work |
| MJM4675 | MJM1100 $\Delta rtmR::bar attTn7::P_{rtmR}-rtmR-3XFLAG-erm$ | This work |
| MJM4920 | MJM1198 $\Delta rtmR::bar$ | This work |
| MJM5028 | MJM1198 $\Delta rtmR::bar attTn7::erm$ | This work |
| MJM5029 | MJM1198 $\Delta rtmR::bar attTn7::P_{rtmR}-rtmR-3XFLAG-erm$ | This work |
| MJM5737 | MJM1100 $attTn7::erm$ / pM1422 | This work |
| MJM5738 | MJM1198 $\Delta rtmR::bar attTn7::erm$ / pM1422 | This work |
| MJM5739 | MJM1198 $\Delta rtmR::bar attTn7::P_{rtmR}-rtmR-3XFLAG-erm$ / pM1422 | This work |
| MJM5788 | MJM1100 $\Delta binK \Delta rtmR::erm-bar$ | This work |
| MJM5847 | MJM1100 $\Delta binK \Delta rtmR::bar$ | This work |
| MJM5850 | MJM1100 $\Delta binK \Delta rtmR::bar$ / pM1422 | This work |
| MJM6110 | MJM1198 $\Delta rtmR::bar attTn7::P_{rtmR}-rtmR-erm$ | This work |
| MJM6155 | MJM1198 $\Delta rtmR::bar attTn7::P_{rtmR}-rtmR-\Delta TM-53-3XFLAG-erm$ | This work |
| MJM6240 | MJM1198 $\Delta rtmR::bar attTn7::P_{rtmR}-rtmR-\Delta TM-72-3XFLAG-erm$ | This work |
| MJM6265 | MJM1100 *rscS*-3XFLAG | This work |
| MJM6266 | MJM1198 *rscS* *-3XFLAG | This work |
| MJM6267 | MJM1198 *rscS* *-3XFLAG $\Delta$ *rtmR*:: *bar* | This work |
| MJM6269 | MJM1198 *rscS* *-3XFLAG *attTn7*:: *erm* | This work |
| MJM6270 | MJM1198 *rscS* *-3XFLAG $\Delta$ *rtmR*:: *bar attTn7*:: *erm* | This work |
| MJM6271 | MJM1198 *rscS* *-3XFLAG *attTn7*::P<sub>*rtmR*</sub> - *rtmR-erm* | This work |
| MJM6272 | MJM1198 *rscS* *-3XFLAG $\Delta$ *rtmR*:: *bar attTn7*::P<sub>*rtmR*</sub> - *rtmR-erm* | This work |
| MJM6282 | MJM1198 Δ*binK* Δ*rtmR*::*bar* | This work |
| MJM6283 | MJM1198 Δ*rtmR*::*bar attTn7*::P<sub>*rtmR*</sub>-*rtmR-RNPnull* | This work |
| MJM6284 | MJM1198 Δ*rtmR*::*bar attTn7*::P<sub>*rtmR*</sub>-*rtmR-RNPnull*-3XFLAG | This work |
| MJM6305 | MJM1198 Δ*binK* Δ*rtmR*::*bar* / pM1422 | This work |
| MJM6310 | MJM1198 Δ*rtmR*::*bar attTn7*::P<sub>*rtmR*</sub>-*rtmR-RNPnull* / pM1422 | This work |
| MJM6321 | MJM1198 Δ*rtmR*::*bar attTn7*::P<sub>*rtmR*</sub>-*rtmR-erm* / pM1422 | This work |
| MJM6325 | MJM1198 Δ*binK* / pM1422 | This work |
| MJM6326 | MJM1100 Δ*rtmR*::*bar* / pM1422 | This work |
| MJM6327 | MJM1198 $\Delta rtmR::bar$ / pM1422 | This work |
| MJM6435 | MJM1100 $rscS^* \Delta cat$ | This work |
| MJM6436 | MJM1100 $\Delta rtmR::bar rscS^* \Delta cat$ | This work |
| *Escherichia coli* |  |  |
| MJM534 | CC118 $\lambda pir$ / pEVS104 | (68) |
| MJM537 | DH5α $\lambda pir$ | Laboratory Stock |
| MJM637 | S17-1 $\lambda pir$ / pUX-BF13 | (71) |
| MJM658 | BW23474 / pEVS107 | (70) |
| MJM1422 | MJM537 / pM1422 | (20) |
| MJM1431 | $\beta 3914$ | (88) |
| MJM3271 | NEB5α / pEVS79-*erm* | (45) |
| MJM3287 | NEB5α / pHB1 | (8) |
| MJM3478 | $\pi 3813$ / pKV496 | (12) |
| MJM3971 | NEB5α / pUCIDT-Kan-IG-VF_2432 | This work |
| MJM4652 | MJM537 / pJVG16 | This work |
| MJM4653 | MJM537 / pJVG17 | This work |
| MJM4879 | MJM537 / pJVG21 | This work |
| MJM6153 | MJM537 / pHN02 | This work |
| MJM6235 | MJM537 / pJVG52 | This work |
| MJM6251 | MJM537 / pJVG55 | This work |
| MJM6273 | MJM537 / pJVG63 | This work |
| MJM6274 | MJM537 / pJVG64 | This work |
| MJM6432 | MJM537 / pJVG21c | This work |

**TABLE 2.**
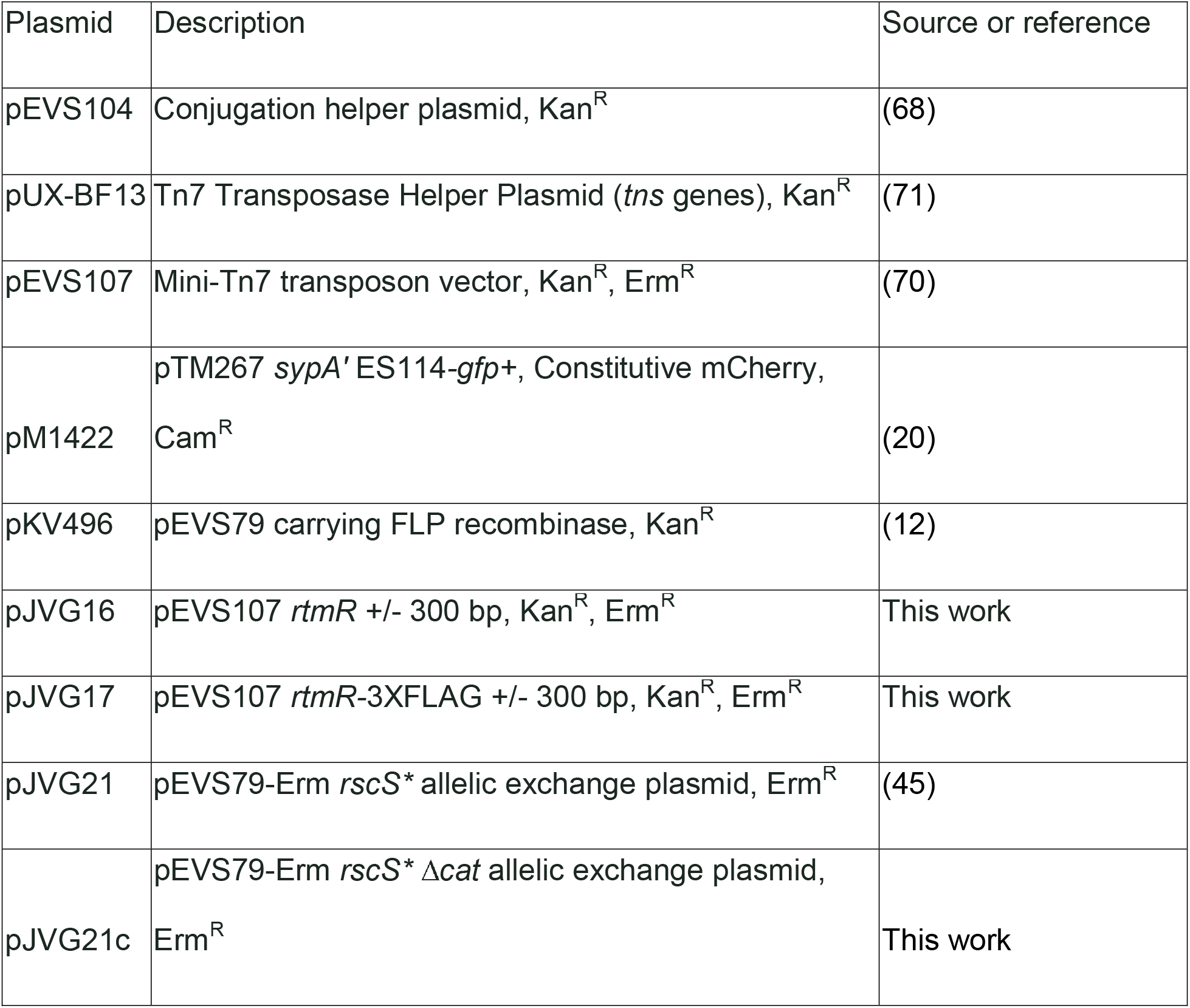
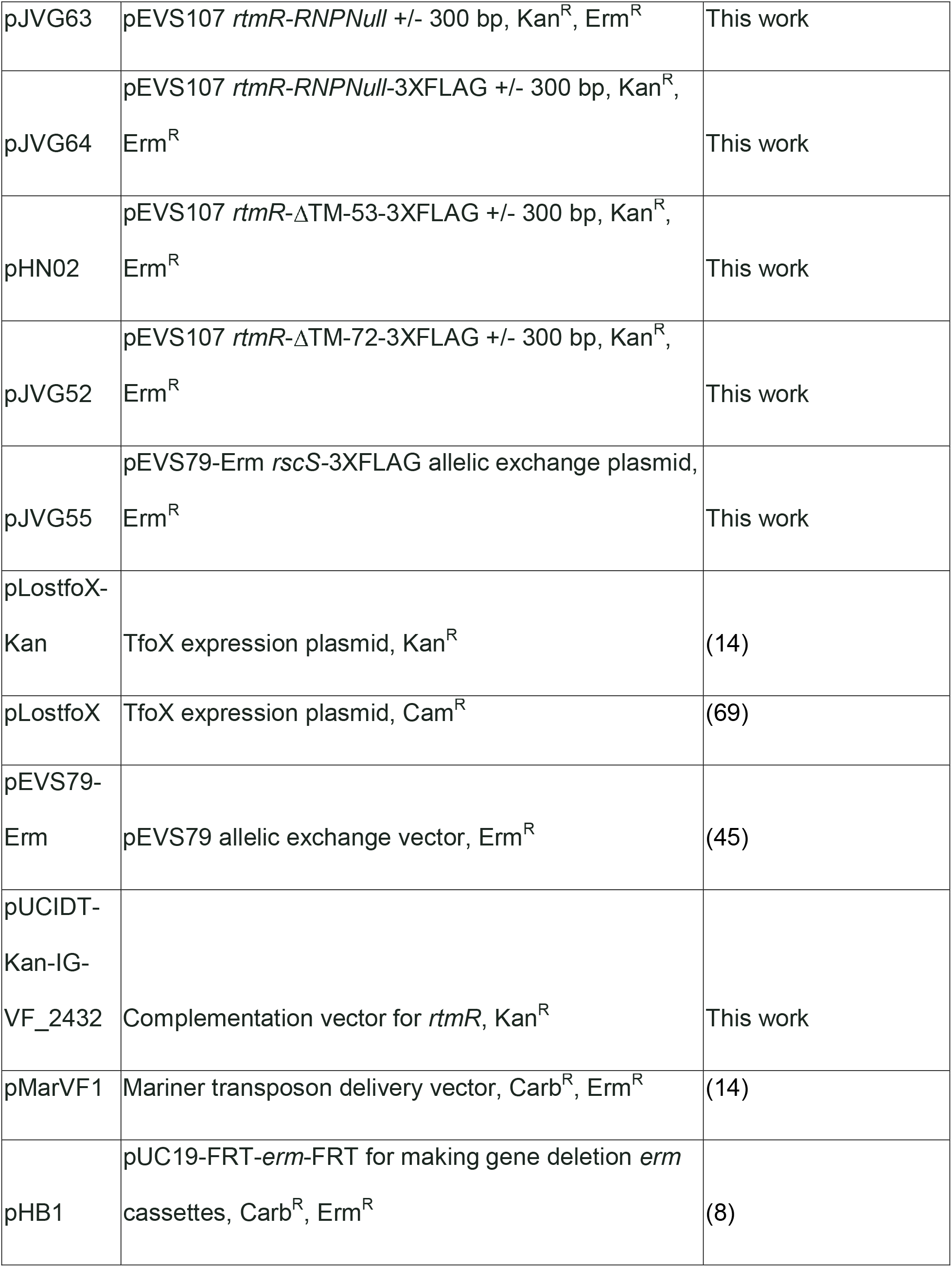
Plasmids used in this work.

### DNA synthesis, purification, and sequencing

Primers employed in this work are listed in **Table S5**. All primers and GeneBlocks were generated by Integrated DNA Technologies (Coralville, IA). Unless otherwise specified, DNA amplicons generated for Sanger and linear amplicon sequencing were amplified using Phusion HS DNA polymerase (NEB), diagnostic PCRs were conducted using GoTaq (Promega) or OneTaq (NEB) DNA polymerases, and DNA amplicons used for cloning were amplified using Q5 DNA polymerase (NEB). All amplicons were purified prior to sequencing, or other usage, using the QIAquick PCR purification kit (Qiagen). All plasmids were purified using the QIAprep spin miniprep kit (Qiagen). When noted, Sanger sequencing was conducted either by the Northwestern University Biotech Center (Chicago, IL) or Functional Biosciences (Madison, WI). Linear amplicon and whole plasmid-sequencing were conducted by Plasmidsaurus (Eugene, OR).

### Transposon mutagenesis of *V. fischeri*

The mariner transposon vector pMarVF1 was conjugated into strain MJM1198 using biparental conjugation with strain MJM1431 (14). Resuspensions of the conjugation spot were plated on LBS-Erm, and allowed to grow at 28 °C for 72 hrs. Colonies showing wrinkled colony morphology at 28 °C were streak purified, and validated for biofilm formation at 25 °C and 28 °C using wrinkled colony spot assays. Transposon insertions were identified using two-round arbitrary PCR. First round amplification was conducted using Arb1 and MJM-440 primers, after which amplicons were purified. Second-round amplification was then conducted on purified amplicons using Arb2 and MJM-477 primers, followed by a further purification step. Amplicons were sent to the Northwestern University Biotech lab for Sanger sequencing (Chicago, Illinois). NCBI BLASTN was used to map Sanger products onto the *V. fischeri* ES114 genome. All hits answering the screen are provided in **Table S6**, yet aside from *binK* and *rtmR* we note that this listing remains to be validated by complementation of the disrupted loci.

### Deletion of *rtmR*

Deletion of *rtmR* was accomplished using a previously published method (13). During the cloning process, we discovered a sequencing error in the ES114 genome (CP000020.2). The incorrect sequence frameshifted *rtmR* after codon 113. The corrected sequence of *rtmR* is provided in **Fig. S10**. All cloning procedures were conducted with the corrected *rtmR* gene, which encodes a predicted 160-amino acid product. The correct nucleotide sequence and gene product of *rtmR* are provided in **Table S7**. Approximately 1 kb upstream and downstream of the *rtmR* CDS was amplified using primers JVG_AC1/2 and JVG_AC5/6 and purified. As *rtmR* is encoded convergently with *murI*, the last 126 bp of *rtmR* were included in the downstream flanking region to prevent off-target effects on *murI*. Amplicons of JVG_AC1/2 and JVG_AC5/6 were mixed 1:1 with an erythromycin resistance cassette amplified from pHB1 using HB_42/154. All three fragments were joined using splicing by overlap extension PCR (SOE-PCR) using the Phusion HS DNA polymerase (NEB). SOE-PCR products were purified, and transformed into *Vibrio fischeri* strain MJM1538 using *tfoX*-mediated natural transformation as performed previously (14, 69). Candidates were screened for Erm resistance, streak purified twice on LBS-Erm, and patched to verify loss of the pLostfoX plasmid. Check PCRs were then conducted to validate insertion of the mutagenic DNA cassette into *rtmR* using JVG_AC1/6, JVG_AC0/HB_08, and JVG_AC3/4. Candidates with predicted disruption of *rtmR* were then amplified using JVG_AC0/7, and sent for Sanger sequencing with primers JVG_AC0, JVG_AC7, HB_08, and HB_09. A passing candidate was then saved as strain MJM4627 (*rtmR::erm-bar*). To remove the erythromycin resistance cassette, plasmid pKV496 carrying the *flp* recombinase was conjugated into MJM4627 using standard triparental conjugation (12). Candidates were streak purified twice on LBS, and patched to ensure the absence of the *erm* cassette (Erm^S^), pKV496 (Kan^S^), along with ensuring no *E. coli* carryover by an absence of growth for LB patches at 37 °C. Check PCR was conducted using JVG_AC0/7 primers, and passing amplicons were sent for Sanger sequencing with JVG_AC0, JVG_AC7, HB_42, and HB_146. A passing candidate was saved as MJM4639 (Δ*rtmR::bar*). This *rtmR* deletion strain is precisely noted as Δ*rtmR::bar* in the methods and strain table and called by the common “Δ*rtmR*” nomenclature in the main manuscript and figures.

### Generation of single-copy complementation alleles of *rtmR*

Complementation of Δ*rtmR* was conducted through Tn*7*-transposon delivered alleles essentially as described previously (45, 70). Plasmid pUCIDT-Kan-IG-VF_2432 was ordered from IDT containing *rtmR* and ∼500 bp upstream (US) and 300 bp downstream (DS) flanking DNA. Primers JVG_BB3/BB12 were used to amplify *rtmR* with ∼300 bp US (P*_rtmR_*) and ∼300 bp DS sequence from plasmid pUCIDT-Kan-IG-VF_2432, and primers JVG_BB10/13 were used to linearize the Tn*7*-transposon vector plasmid pEVS107. The JVG_BB10/13 amplicon was digested with Dpn1 to eliminate template DNA. Amplicons were PCR purified, and joined using NEBuilder HiFi DNA Assembly mastermix (NEB). HiFi products were directly transformed into chemically competent *E. coli* DH5 pir using standard heat-shock treatment. Candidates were streak purified once on LB-Kan, and plasmid DNA was purified. Candidate plasmids were submitted for Sanger sequencing using primers JVG_BB5, 6, 7, and 8, and a passing candidate was saved as MJM4652 (pJVG16). For the 3XFLAG-tagged allele of *rtmR*, the same steps were conducted with the insert DNA instead being amplified from GeneBlock gJVG1, a passing candidate was saved as MJM4653 (pJVG17).

### Design of the RtmR^RNPnull^ allele

The RNP1 and RNP2 motifs present in RtmR were identified through manual annotation of the RRM domain in Benchling, based on previously established consensus motifs (37). Residue swaps capable of disrupting RNA-binding activity within the motifs were identified from the work of Melamed *et al.* (2013) and Deardorff and Sachs (1997) (39, 40). To ensure that the residue changes maintained the proper codon usage levels, each residue to be swapped was matched with the nearest most used codon of the replacement amino acid, determined from the *V. fischeri* ES114 codon usage tables at Kazusa.or.jp. Selected residue swaps, RNP1 - R112Q, F114V, F116V and RNP2 - Y74V, were introduced into the Tn*7* complementation plasmids pJVG16 (Untagged RtmR) and pJVG17 (3XFLAG RtmR) through amplification of either plasmid with primer sets JVG_CL1/4 and JVG_CL2/3. Amplicons of both primer sets were digested with Dpn1 (NEB) to eliminate template DNA, joined with NEBuilder HiFi DNA assembly mastermix, and transformed into chemically competent *E. coli* DH5 pir using standard heat shock procedures. Each candidate was streak purified once on LB-Kan, after which plasmid DNA was isolated and sent for whole plasmid sequencing. Candidates with proper incorporation of the RNP motif residue changes were saved as MJM6273 (pJVG63) and MJM6274 (pJVG64).

### Design of the *rscS*-3XFLAG allele

An allelic exchange vector for inserting a 3XFLAG tag at the end of *rscS* was generated using the pEVS79-*erm* plasmid. In short, the pEVS79-*erm* backbone was amplified using primers RYI097_F/R and homology arms flanking the 3XFLAG insertion site in *rscS* were amplified using JVG_CH1/2 and JVG_CH3/4 from *V. fischeri* gDNA. The RYI097_F/R amplicon was digested with Dpn1 to eliminate template DNA. All fragments including Geneblock gJVG3 were joined using NEBuilder HiFi DNA Assembly mastermix (NEB), and the products were transformed into chemically competent *E. coli* DH5 pir using standard heat shock procedures. Candidates were streak purified on BHI-Erm media once, and overnight cultures from streak purified colonies were saved. Candidates were screened with PCR using primers M13 F/R and sent for whole plasmid sequencing, with a passing candidate saved as MJM6251 (pJVG55).

### Insertion of *rscS-*3XFLAG into *V. fischeri*

Transfer of the *rscS-*3XFLAG allele into strains MJM1100 (Wild type), MJM1198 (*rscS\**), and MJM4920 (*rscS\** Δ*rtmR::bar*) was completed through allelic exchange of plasmid pJVG55 into the chromosome of either strain as described previously (45). The final check PCR for both mutants was conducted with primers JVG_BS6/7, and amplicons were sent for linear amplicon sequencing to validate the presence of the *rscS-*3XFLAG allele. Passing candidates were saved as MJM6265 (*rscS*-3XFLAG), MJM6266 (*rscS\**-3XFLAG), and MJM6267 (*rscS\**-3XFLAG Δ*rtmR::bar*).

### Design of transmembrane truncation mutants

Truncation of the transmembrane helices of RtmR was conducted using the Q5 site-directed mutagenesis kit (NEB). In brief, pJVG17 was amplified with either HN_01/2 (ΔTM-53) or JVG_CB1/3 (ΔTM-72) using Q5 polymerase (NEB), and amplicons were treated with the KLD enzyme mix (NEB). KLD output was directly transformed into *E.coli* DH5 pir, and individual colonies were streak purified on LB-Kan. Candidates were screened for proper assemblies using JVG_AC4/JVG_BK14 (ΔTM-53) or JVG_BB5/6 (ΔTM-72). Candidates passing check PCRs were miniprepped and validated by whole-plasmid sequencing. Sequence-confirmed ΔTM-53 was saved as MJM6153 (pHN02) and ΔTM-72 was saved as MJM6235 (pJVG52).

### Construction of the Δ*binK/rtmR* mutant

Construction of Δ*binK/rtmR* was conducted through TfoX-mediated natural transformation as previously described (13, 14, 69). In short, strain MJM2252 (Δ*binK* pLostfoX-Kan) was selected as the recipient strain, and provided 2.5 µg of gDNA extracted from strain MJM4627 (Δ*rtmR*::*erm-bar*) using the DNeasy Blood and Tissue kit (Qiagen). Candidates that demonstrated erythromycin resistance on LBS-Erm media were streaked twice to purity on LBS-Erm, and screened for insertion of *rtmR*::*erm*-bar and retention of the Δ*binK* mutation using check PCR with primers JVG_AC1/6, JVG_AC0/HB_08, JVG_AC3/4, and DAT11/12. A further check PCR was conducted with primers JVG_AC0/7 flanking *rtmR* and these amplicons were tested with linear amplicon sequencing, resulting in strain MJM5788 (Δ*binK* Δ*rtmR::erm-bar*). The *erm* cassette in MJM5788 was removed through conjugation of the flippase plasmid pKV496 (12) into MJM5788, as described for the creation of the original *rtmR*::*bar* mutant. Passing candidates were screened with PCR using primers JVG_AC0/7 and DAT11/12, and the JVG_AC0/7 amplicons were sent for linear amplicon sequencing to confirm the loss of the *erm* cassette. A sequence-validated candidate was saved as MJM5847 (Δ*binK* Δ*rtmR::bar*).

### Insertion of *rscS\** into the Δ*rtmR* and Δ*binK/rtmR* mutants

Transfer of the *rscS\** allele in strains MJM4639 (Δ*rtmR::bar*) and MJM5847(Δ*binK* Δ*rtmR::bar*) was completed through allelic exchange of plasmid pJVG21 into the chromosome of either strain as described previously (45). The final check PCR for both mutants was conducted with primers JVG_BJ9/10, and amplicons were sent for linear amplicon sequencing to validate the presence of the *rscS\** allele. Passing candidates were saved as MJM4920 (*rscS\** Δ*rtmR::bar*) and MJM6282 (*rscS\** Δ*binK* Δ*rtmR::bar*).

### Tn*7* transposition of pEVS107 backbone plasmids

All Tn*7*-transposon complementation and vector controls generated from pEVS107 were inserted into the *V. fischeri att*Tn*7* site using standard quadriparental conjugation, aided by helper strains MJM534 and MJM637 (71). Candidates demonstrating erythromycin resistance were streaked to purity twice on LBS-Erm, and patched to ensure the loss of helper plasmids and the absence of *E. coli* contamination. Check PCRs were conducted using the Tn*7* Site F and R primers, and the insertion of the Tn*7* transposon was confirmed using Sanger sequencing or linear amplicon sequencing for every strain.

### Wrinkled colony assays

Strains used for wrinkled colony biofilm assays were streaked on LBS, and single colonies were used to inoculate overnight cultures in LBS for 16-18 hrs. The following morning 8 uL of each overnight culture was spotted on LBS agar plates and allowed to grow at the indicated temperatures. Plates were incubated within plastic tupperware containers to maintain consistent humidity. At 24 and 48 hrs post-inoculation, each colony spot was imaged using a Leica M60 microscope equipped with Leica DFC295 camera.

### Fluorescence reporter assays of *sypA* expression

Strains carrying the *sypA’-gfp+* reporter plasmid (pM1422) (20) were streaked on LBS-Cam agar to single colonies. For each biological replicate, a single colony was inoculated into LBS-Cam, and allowed to grow for 16-18 hrs at 25 °C. From each overnight culture 8 uL was then spotted in technical duplicate on LBS-Cam, or LBS-Cam-10 mM Ca^2+^ when specified. Plates were then incubated at the stated temperatures, utilizing the same tupperware containers as in the wrinkled colony assays. After 48 hrs of growth, each colony spot was imaged using a Zeiss Axio.Zoom v.16 fluorescence macroscope for eGFP (488 nm excitation, 509 nm emission, 300 ms exposure time) and mCherry (558 nm excitation, 583 nm emission, 700 ms exposure time) signal, with output quantified by the Zen Blue software suite. Normalized *sypA* expression was calculated as eGFP signal / mCherry signal, with eGFP driven by *sypA’* and mCherry expressed by a constitutive promoter on pM1422. Statistical analyses were completed in Prism 10.

### Quantitative western blotting analyses

Each strain was streaked on LBS and grown overnight at 25 °C. Single colonies were used to inoculate LBS broth cultures grown overnight for 16-18 hrs at the indicated temperatures, and 1 mL of each overnight culture was centrifuged at 8,000 x *g* for 3 min. Supernatants were removed, and the pellet was washed twice with 1 mL 1X TBS (20 mM Tris, 150 mM NaCl, pH: 7.6). Pellets were resuspended in 400 uL of Lysis solution [1% SDS, 1X cOmplete Protease Inhibitor Cocktail (Roche)], mixed thoroughly by pipetting, then 2 uL of TURBO DNAse (Invitrogen) was added. Lysates were incubated for 10 min at 37 °C with 1100 RPM shaking using a ThermoMixer (Thermo Scientific). Lysates were then centrifuged at 20,000 x *g* for 5 min, and the supernatant was removed to a fresh tube. A further centrifugation at 20,000 x *g* for 5 min was used to remove any remaining cellular detritus, and the supernatant was transferred to a fresh tube. Lysates were then frozen at −80°C, with diluted aliquots used to quantify lysate protein concentrations using the DCII kit (BioRad), using bovine serum albumin (BSA) as the standard. On the day of immunoblotting, each sample was thawed, and diluted to a standardized concentration in fresh Lysis solution. Prior to loading, Laemmli sample buffer (Bio-Rad) was added to all samples to a final concentration of 1X with 355 mM 2-mercaptoethanol, and samples were heated at 95 °C for 15 min. Following heating, samples were immediately loaded into 4-20% Mini-PROTEAN TGX precast gels (Bio-Rad), within a mini-cell apparatus filled with 1X Tris/Glycine/SDS buffer [25 mM Tris, 192 mM glycine, 0.1% SDS, pH 8.3] (Bio-Rad). Gels were run at 85V for 15 min, then 200 V until the loading dye reached the bottom of the gel cassette. Following electrophoresis, gels were immediately transferred into bins of 1X Towbin buffer [20% Methanol, 25 mM Tris, 192 mM glycine, pH 8.3] (BioRad), and allowed to rest for 15 min. Gels were transferred to 0.45 µm Immun-Blot Low-Fluorescence PVDF membranes (Bio-Rad) for 30 min at 100 V. After transfer, membranes were dried for a minimum of 1 hr, before staining with the Revert 700 total protein stain kit (LI-COR). Stained membranes were visualized on a LI-COR Odyssey FC imager, and blocked overnight in a blocking buffer [5% non-fat milk, 0.1% Tween-20, 20 mM Tris, 150 mM NaCl, pH: 7.6]. The next morning, each blot was treated with 1:5000 diluted 1 mg/mL mouse M2 IgG anti-FLAG monoclonal antibody (Sigma-Aldrich) in 1X TBS-T [0.1% Tween-20, 20 mM Tris, 150 mM NaCl, pH: 7.6], and incubated with rocking for 1 hr. Membranes were washed 4 times for 15 min with 50 mL 1X TBS-T, then treated with 1:5000 1 mg/ml goat anti-mouse IgG IRDye 800CW (LI-COR) in 1X TBS-T with 0.1% SDS, and incubated with rocking for 1 hr. Following incubation, membranes were washed 3 times with 50 mL 1X TBS-T for 15 min before a final 15 min 50 mL 1X TBS wash. Blots were imaged on a LI-COR Odyssey FC imager, and analyzed for total protein and target signal using ImageStudio (LI-COR), and to verify no sample saturated the imager. Prism 10 was used for statistical analysis.

### Immunoblotting of RtmR expression at 25 °C and 28 °C

Strains MJM4674 and MJM4675 were streaked on LBS and allowed to grow overnight at 25 °C. Single colonies were used to inoculate duplicate 3 mL cultures of LBS, one of which was grown at 25 °C and the other 28 °C for 17-18 hrs. The following morning, 1 mL of each culture was centrifuged at 8,000 x *g*, for 3 min at 4 °C and immediately placed in ice. The supernatant was removed and the pellet was washed twice with 1 mL ice cold 1X TBS. Cell pellets were resuspended in 800 uL Lysis solution [1% SDS, 1X cOmplete Protease Inhibitor Cocktail (Roche)], with 4 uL Turbo DNAse. After this point, samples were treated as described for normal quantitative western blotting as described above.

### Cellular fractionation of *V. fischeri*

Cellular fractionation of *V. fischeri* was conducted with modifications from the method of Matsuda *et al* (36). Strains MJM4674 and MJM4675 were cultured overnight in 30 mL LBS for 17 hrs, after which 20 mL of either strain was centrifuged at 3,000 x *g*, 15 min, at 4 °C. The supernatant was removed, and the pellet was resuspended in 2 mL Periplasting buffer [200 mM Tris-HCl pH: 7.5, 20% Sucrose, 1 mM EDTA, 20 mg/mL Lysozyme (ThermoFisher)]. Cell resuspensions were incubated at room temperature for 5 min, after which 2 mL of ice cold sterile DI H_2_O was added to each sample, followed by a 5 min incubation on ice. Samples were centrifuged at 12,000 x *g,* 2 min, at 4 °C. The supernatant was discarded and the pellet was resuspended in 4 mL 50 mM Tris-HCl pH: 7.5. To each sample, 20 uL of Benzonase (Sigma Aldrich) was added and samples were allowed to rest for 5 min at room temperature. After this, samples were placed on ice and sonicated with a Sonic Dismembrator Model 500 (Fisher Scientific) at 30% amplitude for a total of 20 s. After sonication, each sample was centrifuged at 12,000 x *g* for 10 min at 4 °C and the supernatant was transferred to a fresh tube. A further centrifugation was conducted at 12,000 x *g* for 10 min at 4°C to remove any unlysed cells from the supernatant. For both strains, 100 uL of total lysate was collected and frozen at −80 °C as “Total Lysate”. A 3 mL volume of the MJM4675 sample was centrifuged at 100,000 x *g* for 1 hr at 4°C on an Optima Max XP ultracentrifuge (Beckman Coulter) using a TLA-110 rotor. Cytoplasmic fractions were collected as the top 200 uL of the sample, and the remainder of the supernatant was removed. Membrane pellets were washed twice with 1 mL of 50 mM Tris-HCl pH: 7.5, and the wash was discarded. After the wash steps, the pellets were resuspended in 1 mL 50 mM Tris-HCl pH: 7.5, followed by a 1:2 dilution to ensure that the loading of cytoplasmic and membrane fractions would be equivalent. Both fractions were frozen at −80 °C. The presence of RtmR in Total, Cytoplasmic, and Membrane fractions was assessed using immunoblotting. In brief, samples of each fraction were thawed and 15 uL of each sample was directly mixed with Laemmli sample buffer to a final concentration of 1X with 355 mM 2-mercaptoethanol, and samples were heated at 95 °C for 15 min. Samples were run in 4-20% Mini-PROTEAN TGX precast gels (Bio-Rad), and the immunoblotting process was conducted as described above with Revert 700 total protein staining (LI-COR). Blots were probed with primary 1:5000 0.8 mg/mL rabbit IgG anti-FLAG polyclonal antibody (Sigma-Aldrich) and 1:5000 0.5 mg/mL mouse IgG anti-RpoA antibody (BioLegend), and secondary 1:5000 1 mg/mL goat anti-mouse IgG IR-Dye 680RD (LI-COR) and 1:5000 1 mg/mL goat anti-rabbit IgG IR-Dye 800CW (LI-COR) antibodies. Blots were imaged on a LI-COR Odyssey FC imager, and images were processed in ImageStudio (LI-COR).

### Structural modeling and functional annotation of RtmR

Structural modeling of RtmR was conducted with the predicted full-length 160 amino acid protein using Google ColabFold V.1.5.5 (72), and visualized using UCSF ChimeraX-V1.9 (73). Domain annotation was accomplished using InterProScan (74).

### Generation of a protein phylogenetic tree of RtmR homologs

To discover putative homologs of RtmR, the entire amino acid sequence was used as a query in a TBLASTN search. To ensure that the sequence for each homolog was fully represented, species with hits were identified and whole genomes were downloaded from NCBI for annotation with Prokka (V 1.14.6) (75). Genome annotations were visualized in Benchling, and specific CDS were selected based on TBLASTN hit regions. Predicted homologs were then submitted to the InterProScan web server for functional annotation to confirm the presence of RtmR domain features (74). Validated homologs of RtmR were then aligned in the MEGA11 software suite using the Muscle Aligner using default settings. The JTT+G+I maximum likelihood model was identified by MEGA11 as the best fit to the alignments provided, and thus was used to generate the phylogenetic tree with 250 bootstrap replications (76). NCBI BLASTP was used to determine percent ID to ES114 RtmR.

### Transcriptomics analysis of *rtmR* mutants

Strains MJM3381, MJM5028, MJM5029, and MJM5250 were streaked on LBS and grown overnight at 25 °C. Single colonies of each strain were inoculated into 3 mL LBS and grown for 16 hrs at 28 °C with shaking. Overnight cultures were diluted 1/100 into 20 mL fresh LBS and grown at 28 °C with shaking. At OD600 1.8, 1 mL samples of each strain were taken and centrifuged at 10,000 x *g*, for 1 min. Supernatants were discarded and each pellet was resuspended in 600 uL TriReagent (Zymo Research), and total RNA was purified using a Direct-Zol kit (Zymo Research). Four RNA samples for each strain were submitted to SeqCoast Genomics for sequencing. Sequence files were processed with fastp V0.23.4 (77) before further analysis. Fastp-processed sequence files were then aligned to the *V. fischeri* ES114 reference genome ENSEMBL_GCA_000011805 (MJM3381 and MJM5250) or to ENSEMBL_GCA_000011805-*reform* (MJM5028 and MJM5029) using bwa-mem V0.7.18 (78). For both genomes, information for the 10 currently known sRNAs (VF_2577 *csrB2*, VF_2578 *ryhB*, VF_2579 *spf*, VF_2593 *csrB1*, VF_2597 *qrr1*, VF_2599 *ffs*, VF_2639 *ssrA*, VF_2651 *ssrS*, VF_2654 *rnpB*, VF_A1194 *psrN*) was manually appended. ENSEMBL_GCA_000011805-*reform* is a variant genome that accounts for a chromosomal Tn5 insertion in certain *V. fischeri* strains (referred to as the *rscS\** allele). ENSEMBL_GCA_000011805-*reform* was produced from the genome and annotation files of ENSEMBL_GCA_000011805, and linear amplicon sequencing results of the *rscS\** alle, using the Python-based tool reform (https://github.com/gencorefacility/reform). Reads were assigned to gene features using htseq-count V0.6.1 (79). Differential expression analyses were conducted with EdgeR V4.8.2 (80). One MJM5028 sample was excluded from differential expression analysis as an outlier based on EdgeR MDS plots. All further graphs and data analyses were conducted using RStudio using ggplot2 (81).

### Assessment of anti-FLAG monoclonal antibody specificity

Verification of RtmR-3XFLAG IP with the mouse M2 IgG anti-FLAG monoclonal antibody was conducted essentially as described in Melamed *et al* (2018) (82). Strains MJM4674 and MJM4675 were streaked on LBS and grown overnight at 25 °C. From single colonies of each strain, one 3 mL overnight culture of MJM4674 and two duplicate cultures of MJM4675 were grown overnight at 28 °C. The following morning, each culture was diluted 1/100 in LBS. Growth was tracked until both strains were approximately OD600 0.5, after which culture volumes (in mL) equivalent to (40/OD600) were harvested and centrifuged at 4,500 RPM for 15 min at 4 °C. Supernatants were discarded and pellets were resuspended in 20 mL of ice cold 1X marine phosphate-buffered saline (mPBS) [50 mM Sodium Phosphate, 0.45 M NaCl, pH: 7.4]. Cell resuspensions were re-centrifuged at 4,500 x *g* for 10 min at 4 °C. Supernatants were discarded, and the wash step was repeated. The washed cell pellet was resuspended in 10 mL 1X mPBS and subjected to 80 mJ/cm^2^ UV irradiation (254 nm) on a HL-2000 HybriLinker (Fisher Scientific). After UV exposure, samples were centrifuged at 4,500 x *g* for 10 min at 4 °C. The supernatant was discarded and the pellet was resuspended in 1 mL 1X mPBS, followed by centrifugation at 17,000 x *g* for 3 min at 4 °C. The supernatant was removed, and the cell pellet was frozen in liquid N_2_ and stored at −80 °C until lysis. On the day of lysis, cell pellets were thawed on ice and resuspended in 800 uL ice cold wash buffer [49.2 mM Sodium Phosphate, 295 mM NaCl, 10 mM Imidazole, 0.1% IGEPAL] supplemented with EDTA-Free Protease Inhibitor Cocktail Set III (Calbiochem). Cell resuspensions were transferred to 2 ml screwcap tubes filled with 400 uL of 0.1 mm glass beads (Cole-Parmer). Cells were lysed through vortexing at max-speed on a Vortex-Genie 2 equipped with a MN bead tube holder (Macherey-Nagel) for 2.5 min at 4 °C, after which tubes were rested on ice for 2 min. This bead beating and cooling cycle was repeated 3X. After bead-beating, tubes were centrifuged at 17,000 x *g* for 2 min at 4 °C. Approximately 600 uL of supernatant was transferred to a 1.7 mL MAXYMum recovery tube (Sigma). A further 400 uL of cold wash buffer was added to the lysate remaining on the beads, and the bead-beating and cooling cycle was repeated a further 2X. The beads were again centrifuged at 17,000 x *g* for 2 min at 4 °C, and the supernatant was added to the lysate MAXYMum recovery tube. This combined lysate was centrifuged at 17,000 x *g* for 15 min at 4 °C to remove any unlysed cells. The lysate was transferred to a fresh MAXYMum recovery tube, and 24 uL of the total lysate of MJM4674, and the two duplicate MJM4675 cultures were saved as total lysate samples with 8 uL of 4X Laemmli Dye (Bio-Rad) and frozen at −20°C. In preparation for the immunoprecipitation, 20 uL of Protein A/G magnetic beads (Pierce) were washed with 200 uL of cold wash buffer, before the addition of 750 uL of lysate for pre-clearing. Lysates were then rotated for 60 min at 4 °C on a Hulamixer (Life Technologies). Following lysate pre-clearing, samples were transferred to tubes with 20 uL of Protein A/G magnetic beads loaded with 3 uL of 1 mg/mL mouse M2 IgG anti-FLAG monoclonal antibody for strains MJM4674, and one of the MJM4675 lysates. As a negative control, one MJM4675 lysate was immunoprecipitated with 20 uL of unloaded Protein A/G magnetic beads. Samples were allowed to immunoprecipitate for 90 min with rocking on a Hulamixer at 4 °C. At the completion of the immunoprecipitation, beads were precipitated on a magnetic rack and 24 uL of supernatant was taken as the unbound lysate into 8 uL 4X Laemmli dye and frozen at −20°C. The remaining supernatant was discarded, and the beads were washed for 10 min with 200 uL cold wash buffer, this wash step was repeated 4 times. The final washed beads were resuspended in 15 uL 4X Laemmli dye, and frozen at −20°C. The following day, all samples were thawed on ice, incubated at 55°C for 5 min, and run in 4-20% Mini-PROTEAN TGX precast gels (Bio-Rad), and the immunoblotting process was conducted as described above with Revert 700 total protein staining (LI-COR). Blots were probed with primary 1:5000 0.8 mg/mL rabbit IgG anti-FLAG polyclonal antibody (Sigma-Aldrich) and 1:5000 0.5 mg/mL mouse IgG anti-RpoA antibody (Biolegend), and secondary 1:5000 1 mg/mL goat anti-mouse IgG IR-Dye 680RD (LI-COR) and 1:5000 1 mg/mL goat anti-rabbit IgG IR-Dye 800CW (LI-COR) antibodies. Blots were imaged on a LI-COR Odyssey FC imager, and images were processed in ImageStudio (LI-COR).

### Preparation of L3-IR-App iiCLIP-seq adaptor

Preparation of the L3-IR-App iiCLIP-seq adapter was conducted as described in Lee *et al* (41).The L3-IR-Phos adaptor was synthesized by IDT, and resuspended in ultrapure nuclease-free H_2_O to a final concentration of 100 µM. The adapter was adenylated with the 5’ DNA adenylation kit (NEB), and incubated at 65 °C for 2 hrs, followed by a 10 min 85 °C heat inactivation step. Products were precipitated with 3 M sodium acetate pH 5.5 and ice cold 100% EtOH overnight for 18 hrs at −20 °C. The following morning, the products were centrifuged at 21,000 x *g* for 30 minutes at 4 °C. The supernatant was removed and the pellet was gently washed with ice-cold 80% EtOH. The pellet was then air-dried for 5 minutes and resuspended in 180 uL 1X RNAse-free PBS pH 7.4, and a 1 hr incubation at 37 °C was provided to enable the pellet to dissolve before the dye-tagging step. The adapter was then provided with 20 uL of 10 mM IR-Dye 800CW DBCO dye, and allowed to incubate for 2 hours at 37 °C on the thermomixer. After 2 hrs, the reaction mixture was diluted in 4.8 mL PNI buffer (1:1.5 Qiagen Buffer PB, 100% Isopropanol), and mixed. Aliquots of 250 uL of the product mix were added to Qiagen QIAquick spin columns, and centrifuged for 30s at 6,000 rpm. Flowthrough was discarded, and the columns were washed with 750 uL of 80% EtOH through centrifugation at 13,000 rpm for 1 min. The flowthrough was discarded, and the columns were dried at 13,000 rpm for 2 min. Columns were eluted with 50 uL Ultrapure nuclease free H_2_O for 2 minutes, and centrifuged into clean microtubes at 13,000 rpm for 1 minute. All flowthrough was pooled and quantified using a NanoDrop. The final product was diluted to 1 µM and saved as the L3-IR-App adaptor at −20 °C.

### RtmR iiCLIP

iiCLIP-seq of RtmR was conducted using the method of Lee *et al.* with modifications for *V. fischeri* (41). Single colonies of strain MJM5029 (*rscS\** Δ*rtmR att*Tn*7*::*rtmR*-3XFLAG) were grown for 16 hrs at 28 °C in LBS media with 220 RPM shaking, and the following morning subcultures were diluted 1/100 into 100 mL of fresh LBS media. Subcultures were grown at 28 °C until OD600 1.8, after which cultures were immediately placed on ice. Two 25 mL aliquots of the subculture were poured into sterile Nunc Omnitrays (Fisher Scientific), and subjected to 800 mJ/cm^2^ UV irradiation (254 nm) on a HL-2000 HybriLinker (Fisher Scientific). Both crosslinked aliquots were decanted into a 50 mL conical and centrifuged for 15 min at 6,000 x *g* and 4 °C. Supernatants were discarded and pellets were resuspended in 4 mL ice cold Lysis Buffer [50 mM Tris-HCl pH: 7.4, 100 mM NaCl, 1% IGEPAL CA-630, 0.1% SDS, 0.5% sodium deoxycholate, 1X cOmplete Protease Inhibitor cocktail (Roche)]. Resuspended pellets were mixed 1:1 with 0.1 mM glass beads (Cole-Parmer) in 2 mL screwcap tubes, and vortexed at max-speed on a Vortex-Genie 2 equipped with a MN bead tube holder (Macherey-Nagel) for 2.5 min at 4 °C, after which tubes were rested on ice for 2 min. This bead beating and cooling cycle was repeated 3X. After the final bead beating step, tubes were centrifuged for 2 min at 17,000 x *g* and 4 °C. The supernatants were collected, and lysate protein concentration was determined using the DC II protein kit (BioRad), using BSA to generate the protein standard curve. Lysates were diluted to a final protein concentration of 1 mg/mL in Lysis Buffer, and separated into four separate 1 mL aliquots. RNA ligands were trimmed using RNAse I (ThermoFisher) using either “high” (0.8U) or “low” (0.4U) doses, with both conditions receiving 2 uL of TURBO DNAse (Invitrogen), at 37 °C for 3 min with 1100 RPM shaking on a Thermomixer (Thermo Scientific). After this incubation, samples were cooled for 5 min on ice, before centrifugation at 21,000 x *g* for 10 min at 4 °C. Supernatants were then transferred to Proteus Clarification Mini Spin columns (Anatrace), and centrifuged at 13,000 RPM for 1 min at 4 °C. Final lysate samples were then applied to Dynabeads Protein G magnetic beads (Thermo) loaded with 4 ug of mouse M2 IgG anti-FLAG monoclonal antibody, or non-specific mouse IgG (Santa Cruz), and allowed to rock overnight at 4 °C. The following morning, the beads were dephosphorylated using PNK (NEB) and FastAP Alkaline Phosphatase (ThermoFisher) at 37 °C for 40 min with 1100 RPM shaking. After PNK and FastAP treatment, the 3’ pre-adenylated L3-IR-App adaptor was ligated onto the bead RNA-protein complexes using T4 RNA ligase I (NEB), PNK, and a final adaptor concentration of 0.1 µM. Beads were incubated at room temperature (23 °C) for 75 min, and tubes were flicked every 10 min to ensure mixing of beads with ligation mix. After ligation, unligated adaptors were removed with an enzyme mix of 5’ Deadenylase (NEB) and RecJ_f_ endonuclease (NEB), incubated at 30 °C for 1 hr, then for 30 min at 37 °C, shaking at 1100 RPM. RNA-protein complexes were eluted from beads using 1X NuPAGE loading buffer (Thermo) with 0.1 M DTT, and incubated at 70 °C for 1 min. Elutions were then loaded into 4-12% NuPAGE Bis-Tris polyacrylamide gels (Thermo), and run in 1X NuPAGE MOPS running buffer (Thermo) supplemented with NuPAGE antioxidant (Thermo) at 180V until the loading dye front reached the bottom of the gel, and the dye front was excised before transfer. Gels were then transferred to 0.45 µm Protran nitrocellulose membranes (Amersham) in 1X NuPAGE transfer buffer (Thermo), supplemented with 10% Methanol and NuPAGE Antioxidant, at 10 V for 1 hr. Membranes were then imaged using a LICOR Odyssey FC imager, and immediately frozen at −80 °C. All three complexes were harvested from the anti-FLAG samples treated with the lowest RNase digestion condition (0.4U) in both biological replicates. RNA ligands were liberated from the RtmR complexes using Proteinase K treatment of membrane slices at 50 °C for 1 hr at 1100 RPM on a thermomixer, followed by Phenol:Chloroform extraction. The aqueous fraction was precipitated overnight at −20 °C with sodium acetate pH: 5.2, EtOH, GlycoBlue (ThermoFisher), and 0.5 pmol of a sample-specific reverse transcription primer. The following morning RNA pellets were recovered and reverse-transcribed using SuperScript IV (Invitrogen). RNAs were then degraded with NaOH treatment followed by neutralization with HCl, and cDNA was purified using Agencourt AMPure XP beads (Beckman Coulter). Circularization of cDNA was conducted using CircLigase II (LGC), after which a further bead cleanup was performed. Phusion HF polymerase (Thermo) was used for amplifying cDNA libraries using P3 and P5 Solexa primers with 22 cycles. Final libraries for two biological replicates were purified using bead cleanup, and submitted for sequencing at the University of Minnesota Genomics Center using an Illumina NextSeq 2000 platform.

### Analysis of RtmR iiCLIP Data

Reads for the RtmR iiCLIP-seq output from the NextSeq sequencing run were first demultiplexed using Ultraplex V.1.1.5 (83), using barcodes NNNNTCCACNNN (Rep. 1) and NNNNTTTAANNN (Rep. 2). Next, demultiplexed reads were analyzed using the nf-core/clipseq analysis pipeline V1.0.0 (42) against the *V. fischeri* ES114 ENSEMBL_GCA_000011805-*reform* genome. Information on all unique tRNA and rRNA genes present in *V. fischeri* were provided as a separate file for smRNA pre-mapping. Within the nf-core/clip-seq pipeline, parameters for the STAR mapper were modified as previously described in Monti *et al.* (2024) (84): “--outFilterMultimapNmax 1 --outFilterMultimapScoreRange 1 --outSAMattributes All -- alignIntronMin 1000000 --outFilterScoreMin 10 --alignEndsType Extend5pOfRead1”. We opted to use --outFilterMultimapNmax set to 1 instead of 100 as used previously for more stringent mapping. Peakcaller iCount was invoked, and the adapter sequence “AGATCGGAAGAGCGGTTCAG” was provided. Statistically significant iCount peaks identified using a 3 nt half window were joined using a 3 nt merge window and used for further analysis of RtmR target genes in RStudio. Peaks were then summed by peak score within their respective genes, and plotted using ggplot2 V3.5.2. To visualize the crosslink positions across specific transcripts, clipplotr V1.0.0 (85) was used with the raw crosslink positions output from the nf-core/clip-seq pipeline. For each gene analyzed, 100 bp of flanking sequence was provided on each side of the gene.

### Deletion of *cat*

The plasmid pJVG21 was modified using Q5 site-directed mutagenesis (NEB) to remove the *cat* ORF. Primers DAT_546 and DAT_548 amplified pJVG21 and the amplicon was gel-purified (Qiagen QIAquick gel extraction kit) and treated with the KLD enzyme mix (NEB). This was transformed into *E.coli* DH5 pir, and individual colonies were streak purified on BHI-Erm. Candidates were sent for whole-plasmid sequencing to confirm correct assembly. Using this new plasmid, pJVG21c (strain MJM6432), allelic exchange was conducted in strains MJM1100 (ES114) and MJM4639 (MJM1100 Δ*rtmR*::*bar*), as described previously (45). Cells that incorporated the plasmid began wrinkling and wrinkled colonies were preferentially chosen as candidates during the allelic exchange process. A final check PCR was done with primers DAT_015 and DAT_016. Final candidates were also sent for whole genome sequencing (Plasmidsaurus) to confirm both the *rscS\** and *rtmR* regions of the chromosome had the desired sequence. The validated candidates were saved as MJM6435 and MJM6436.

## ACKNOWLEDGMENTS

We thank Sarah Quillin and David Luy for early work on this project, and Joseph Dillard and Ryan Schaub for providing access and advice regarding the use of the Optima MAX-XP ultracentrifuge. This work was supported by NIGMS R35GM148385 to M.J.M. A.M. is an H.I. Romnes Faculty Fellow funded by the Wisconsin Alumni Research Foundation and a Vilas Faculty Mid-Career Investigator. J.A.V.G. was supported by the NIAID training grant T32AI055397. The content is solely the responsibility of the authors and does not necessarily represent the official views of the National Institutes of Health.

## DATA AVAILABILITY

iiCLIP-seq data were deposited on the Gene Expression Omnibus (GEO) under series accession GSE301959. RNA-seq data were deposited on Gene Expression Omnibus (GEO) under series accession GSE337776.

