## Supplementary figures and images for "RtmR is a membrane-embedded RRM-family RNA-binding protein"

### Figure S1

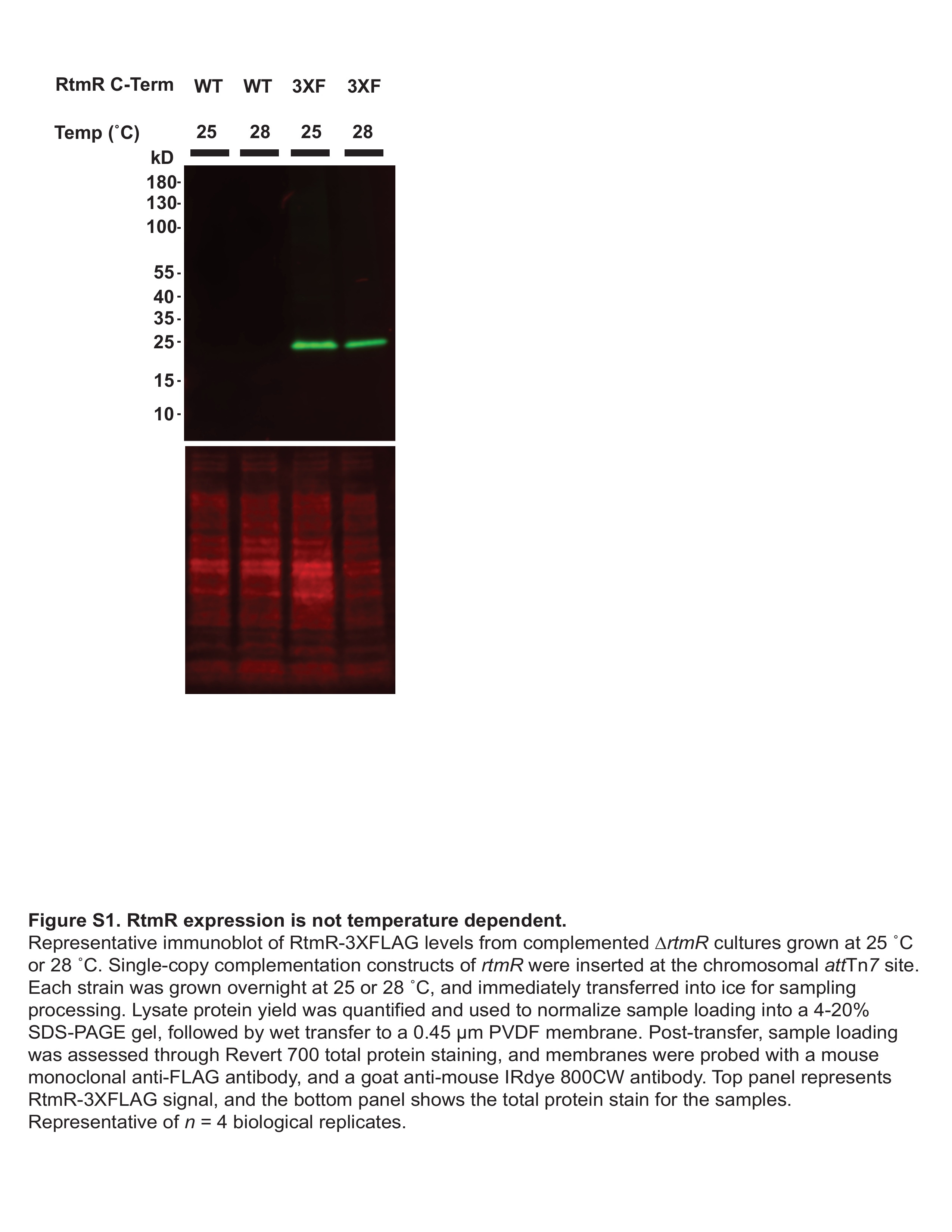

### Figure S2

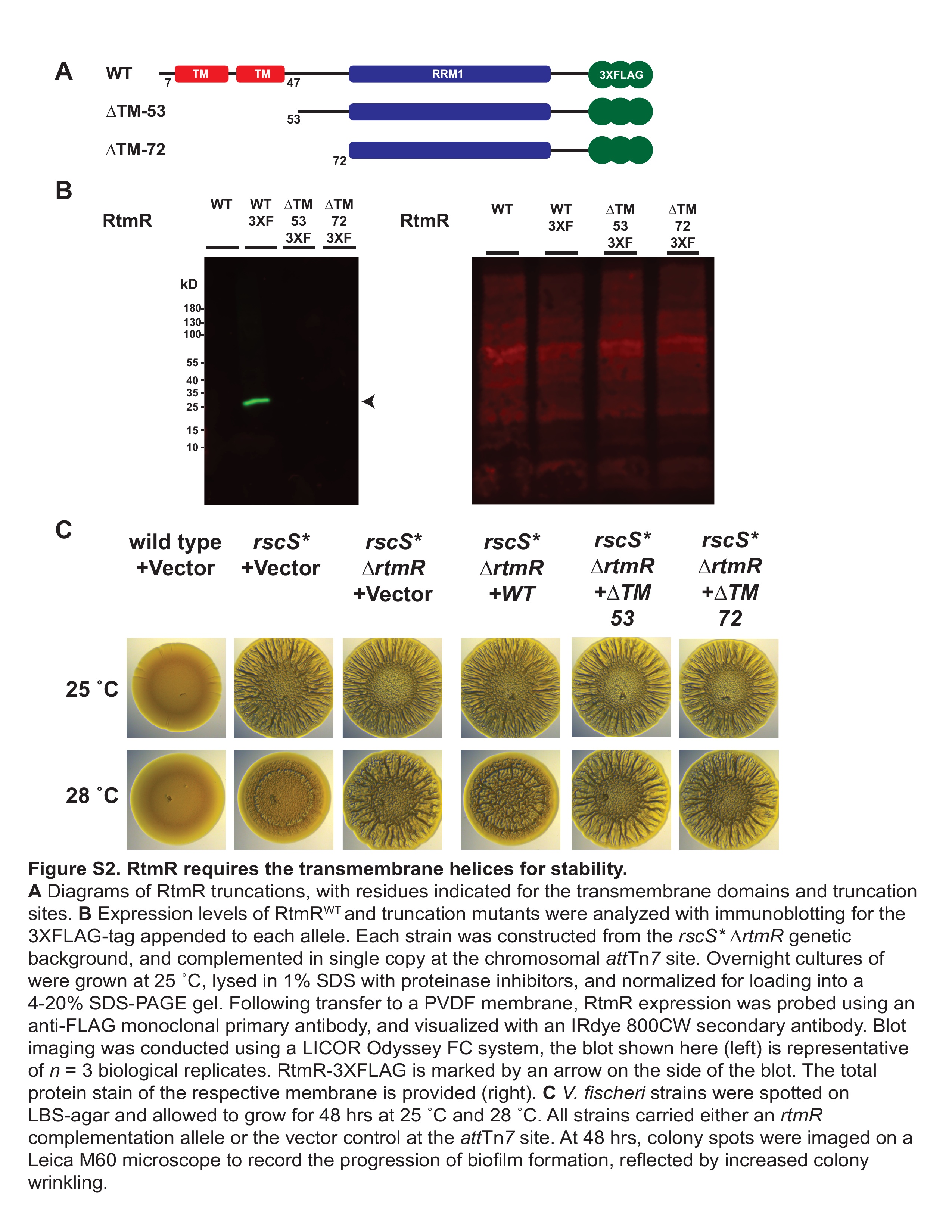

### Figure S3

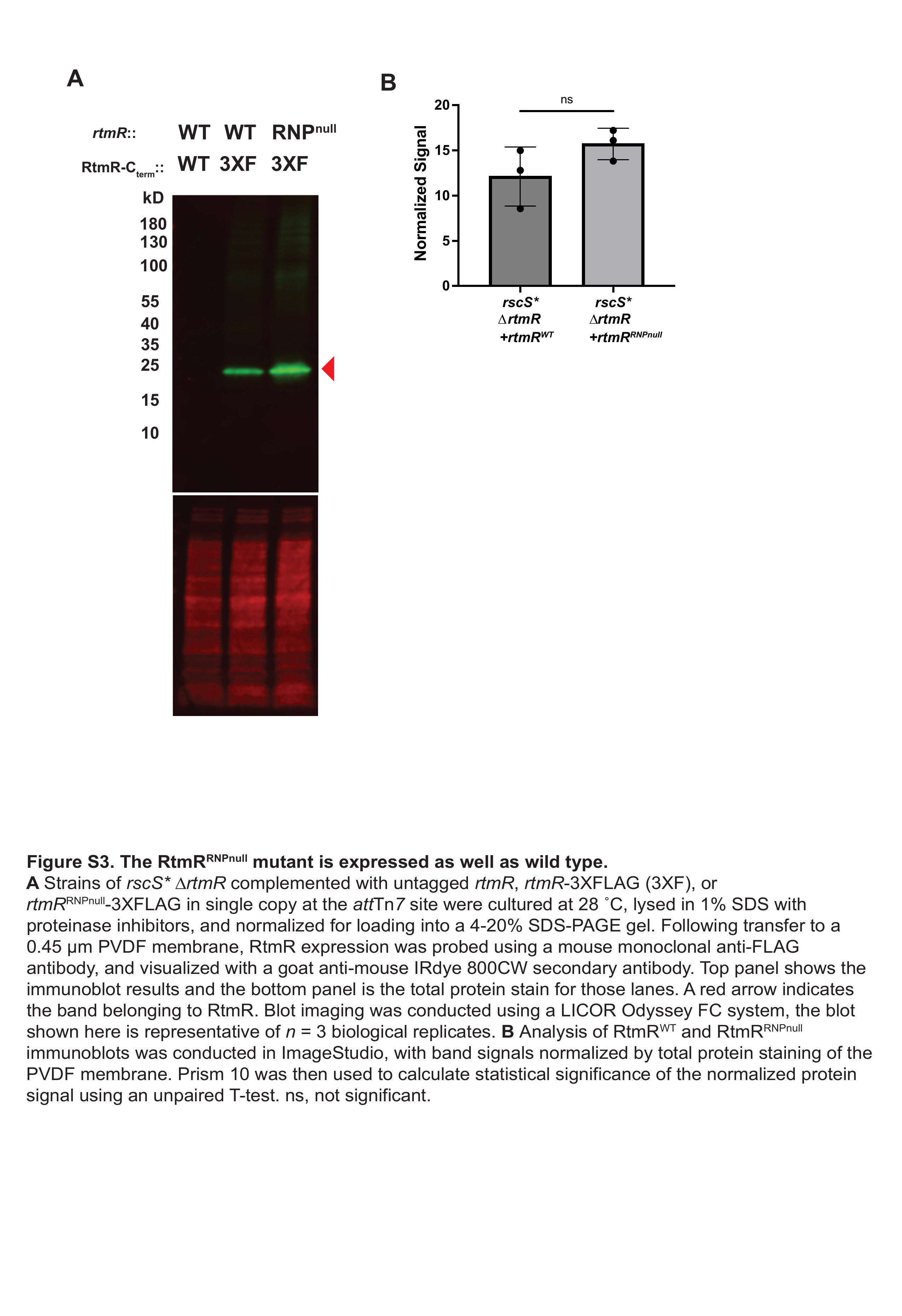

### Figure S4

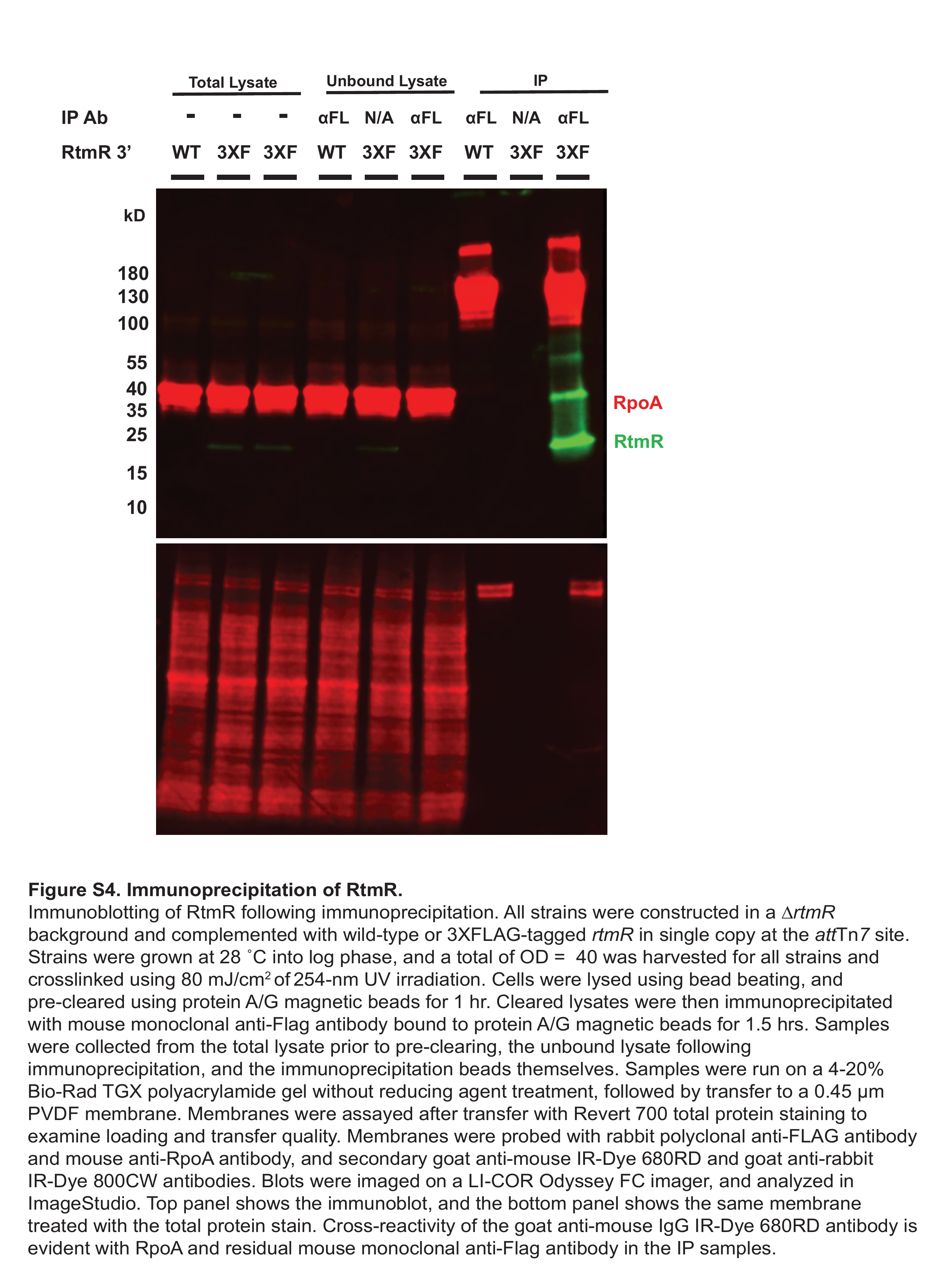

### Figure S5

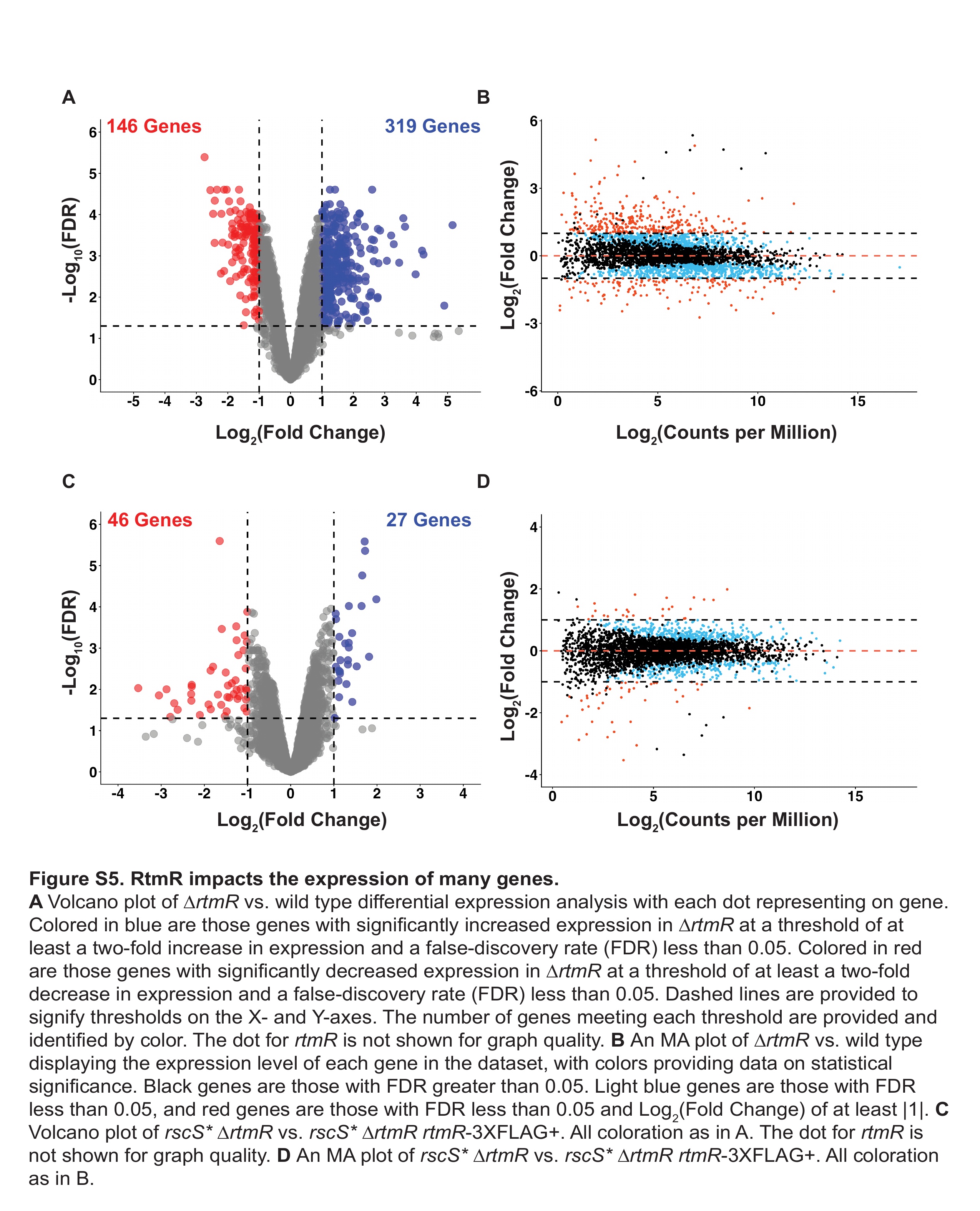

### Figure S6

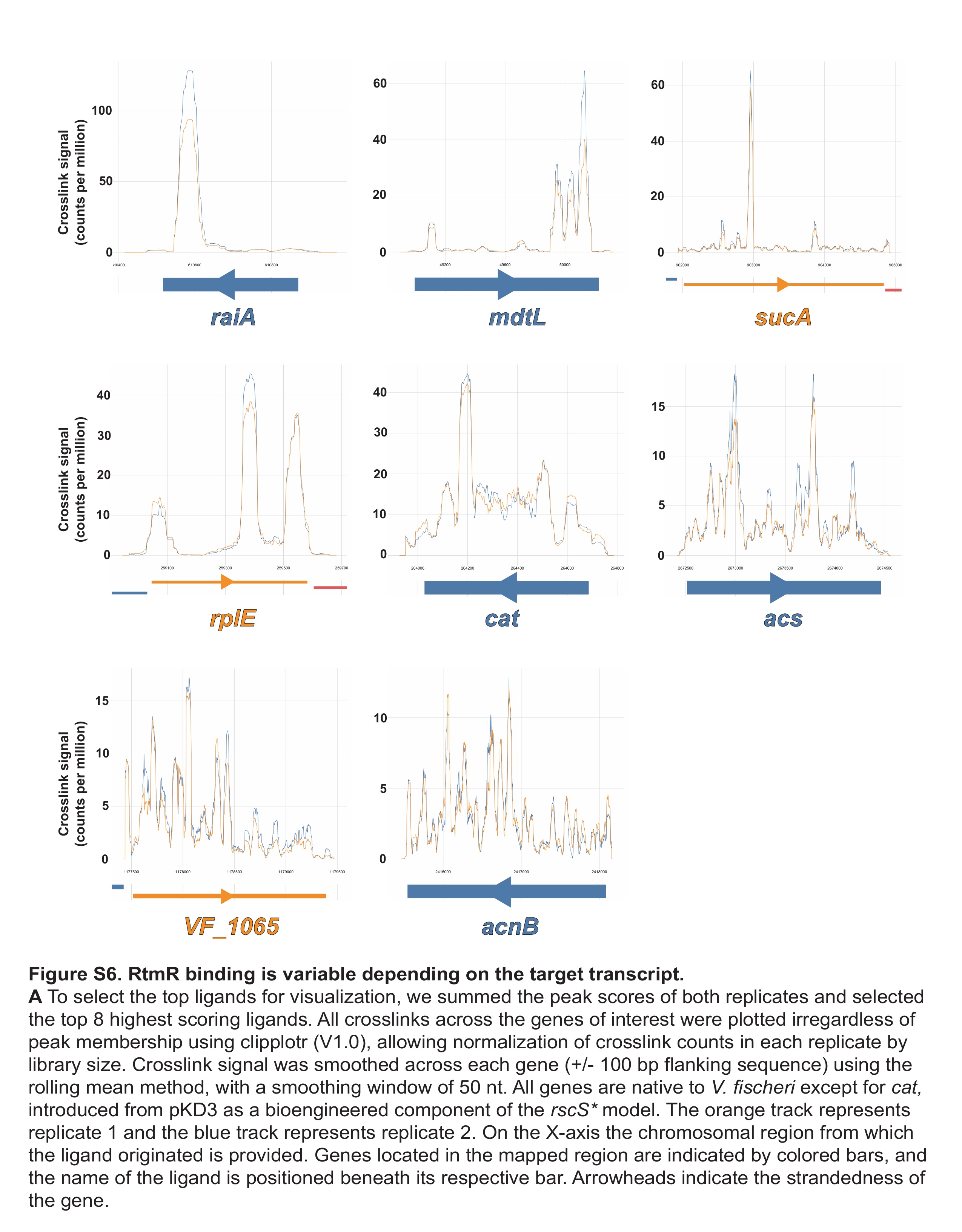

### Figure S7

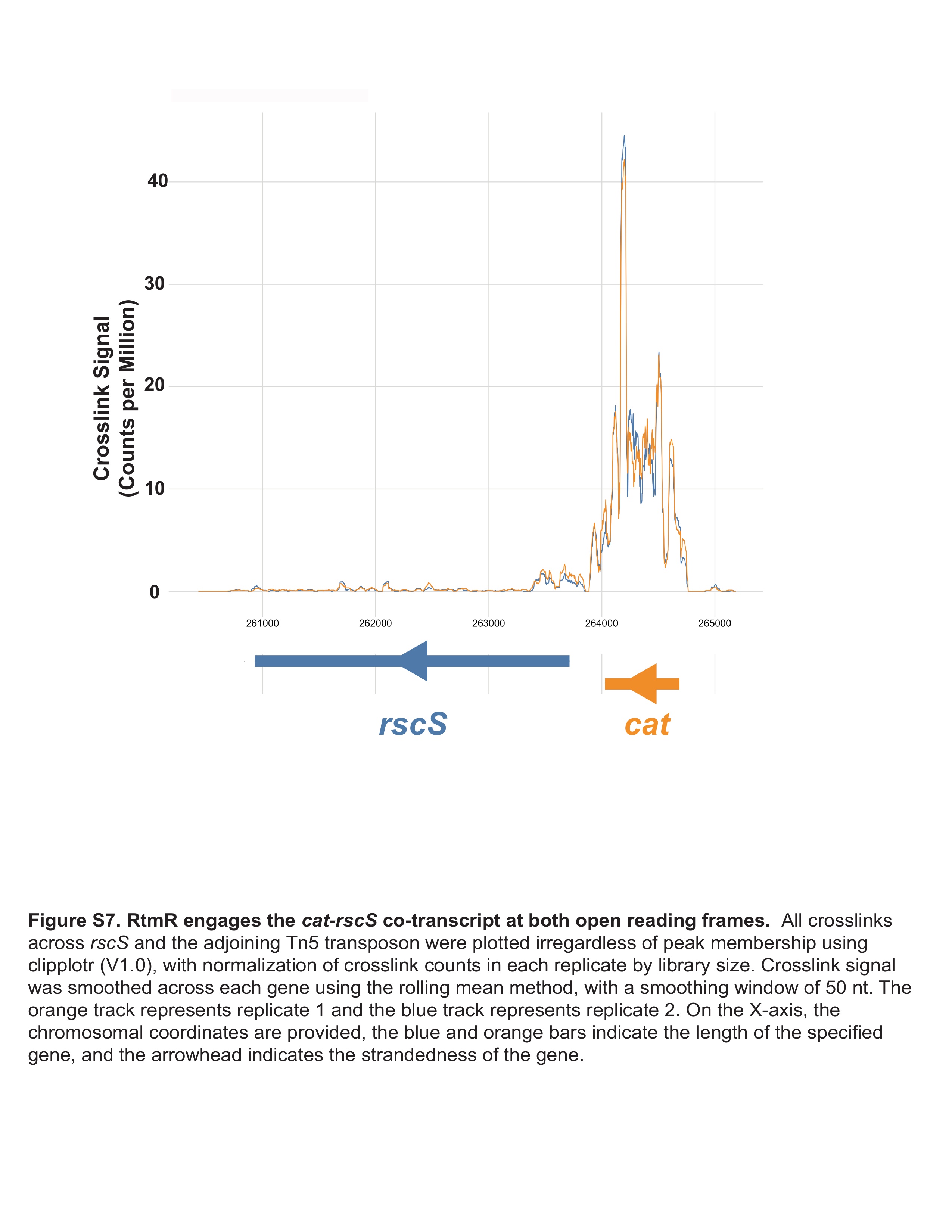

### Figure S8

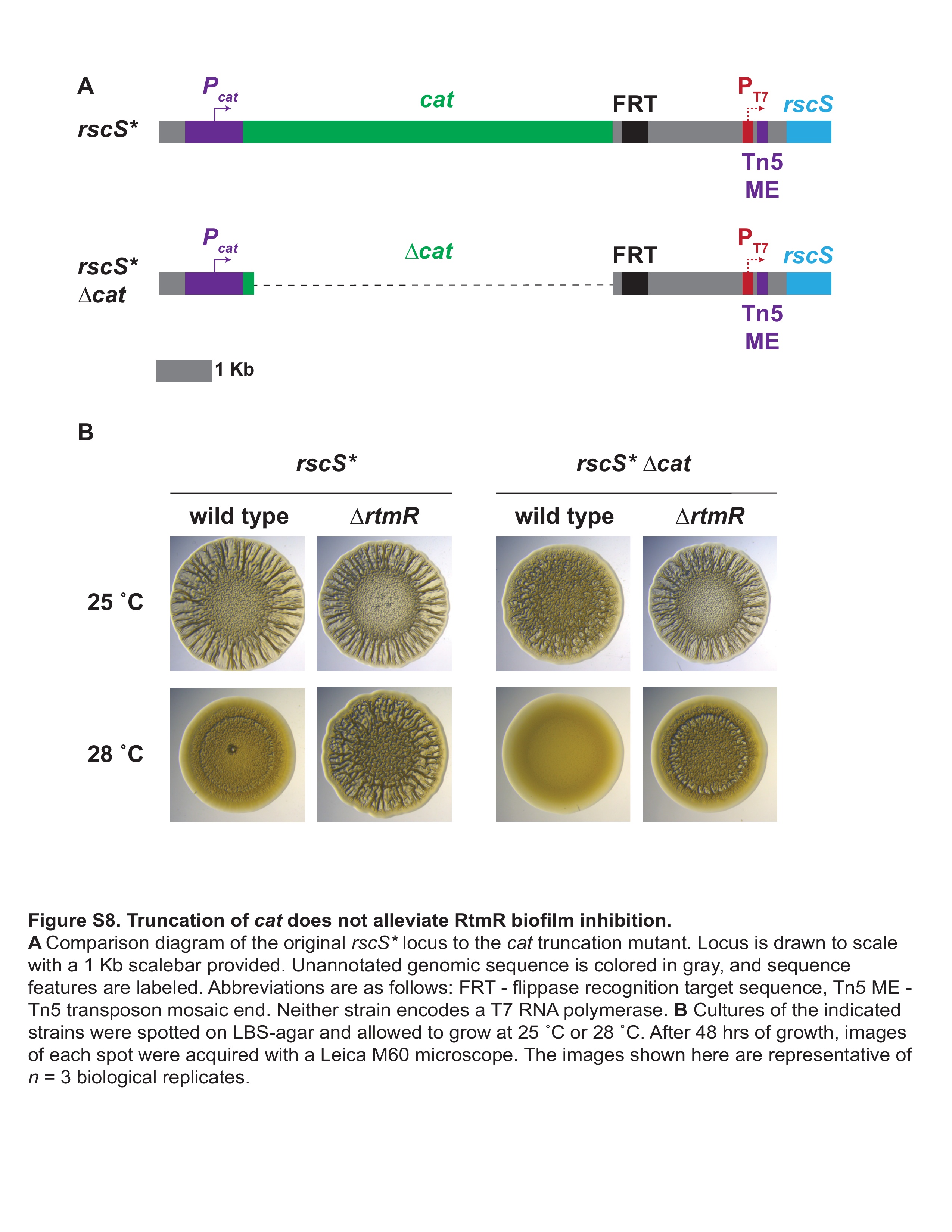

### Figure S9

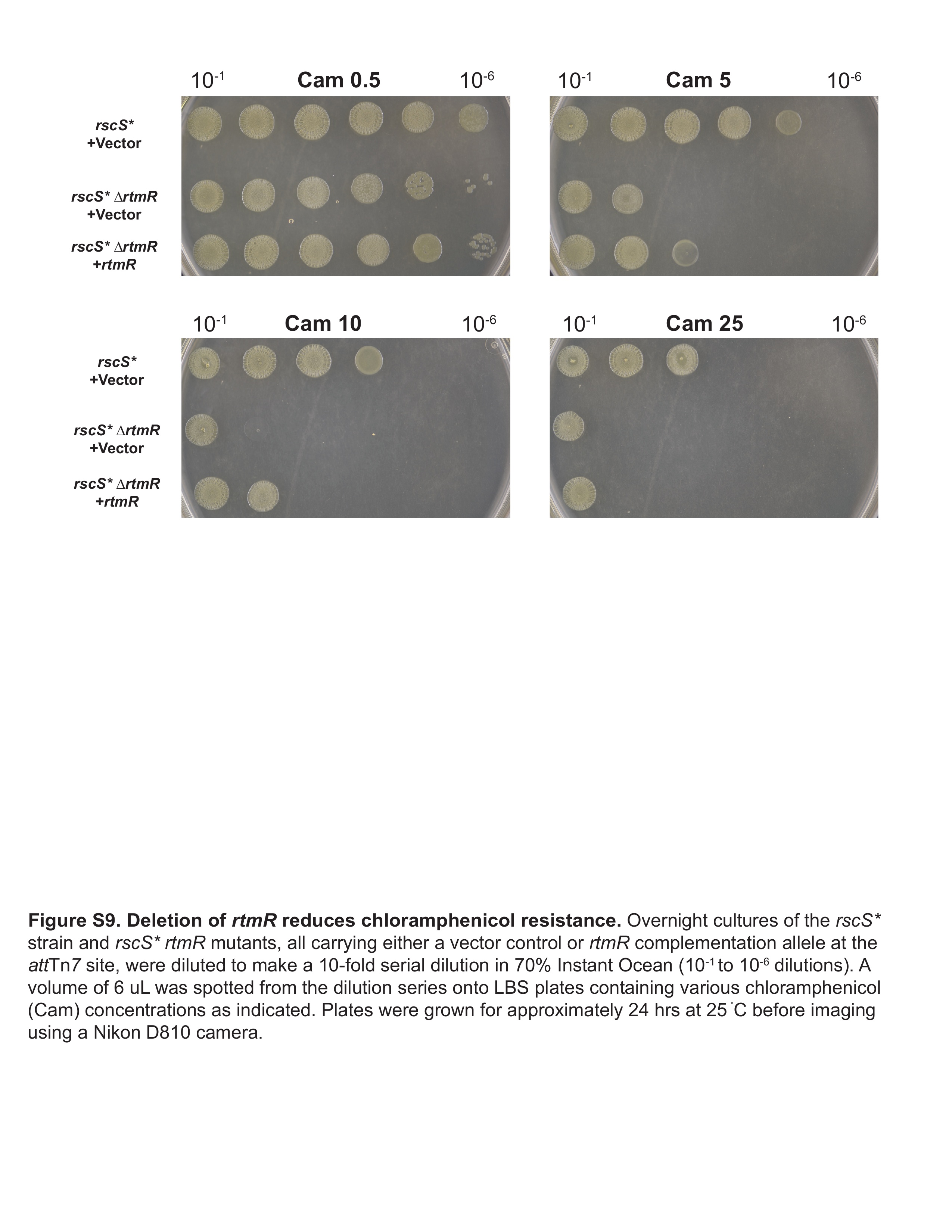

### Figure S10

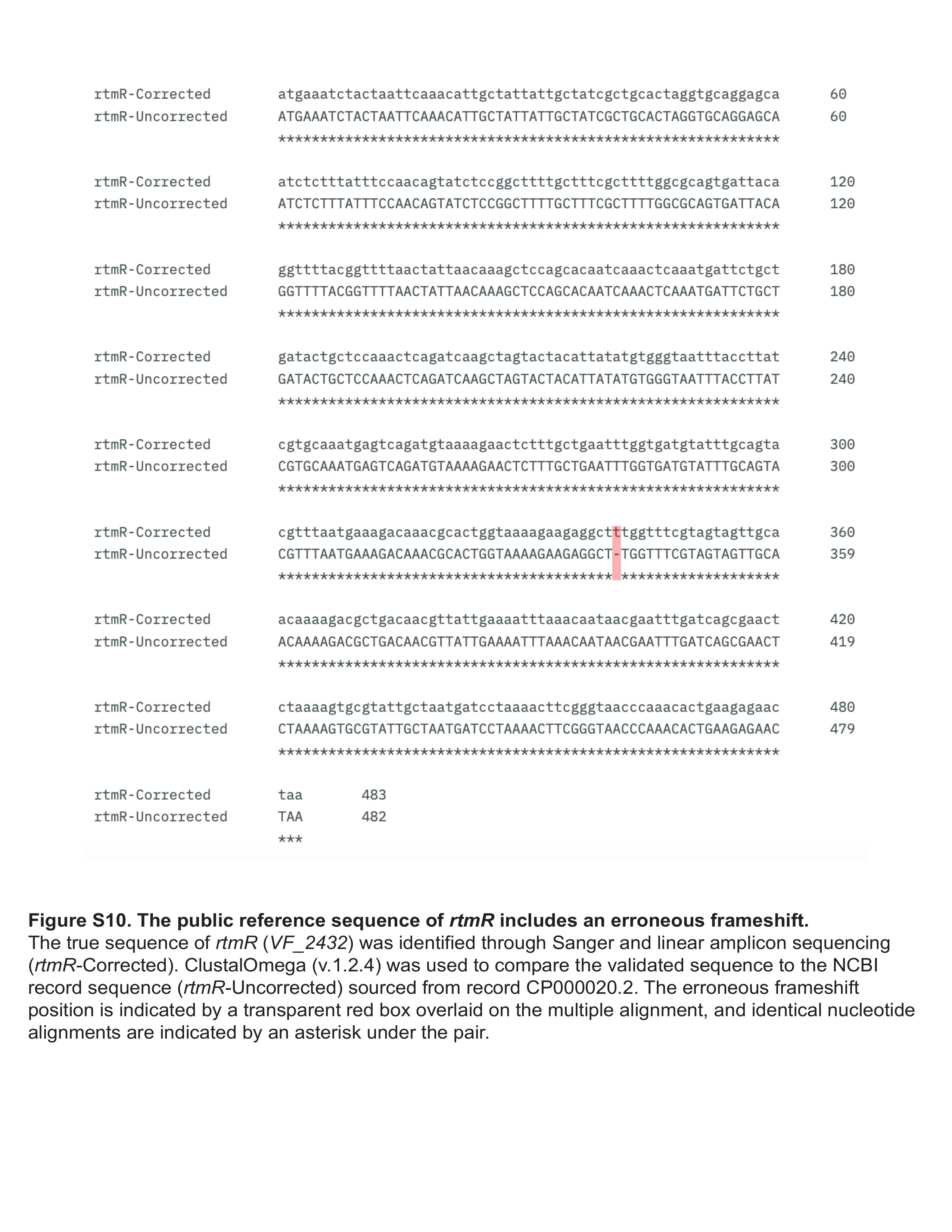
